# Vesicle architecture licenses antigen-specific memory T cell immunity after mucosal immunization with infection-derived vesicles

**DOI:** 10.64898/2026.09.09.750371

**Authors:** Saloni Bhimani, Alexander C. Schultz, Sae Tanaka, Mariola J. Ferraro

## Abstract

Small extracellular vesicles (sEVs) released by infected cells carry microbial antigens together with host immunomodulatory molecules, yet whether their structural integrity governs the durability of protective immunity is unknown. Using *Salmonella* infection as a model, we show that sEVs isolated from infected macrophages act as potent antigen carriers for intranasal immunization. In BALB/c mice, intact sEVs elicited antigen-specific serum IgG responses comparable in magnitude to those induced by live attenuated Δ*aroA Salmonella.* Immunization further increased fecal IgA binding to *Salmonella* following lethal challenge and generated serum capable of enhancing macrophage-mediated bacterial uptake. Single-cell RNA sequencing of mesenteric lymph nodes revealed enrichment of mature B cells, effector CD8 T cells, and central memory–like CD4 T cells expressing CD69, CD44, CCR7, and Sell/CD62L, while defined recombinant sEV-associated antigens drove IFN-γ and IL-2 production by memory T cells ex vivo. Disrupting vesicle architecture by sonication left total IgG and IgG1 induction intact but selectively impaired IgG2a responses and antigen-specific T-cell recall; intact sEVs, by contrast, drove greater dendritic-cell CD80 expression, IL-2 production by cocultured CD69+ CD4 T cells, and prolonged survival after lethal challenge. These findings establish that vesicle integrity is dispensable for the overall magnitude of antibody induction but is required to instruct dendritic-cell costimulation, Th1-associated IgG2a class switching, and durable memory T-cell recall after mucosal immunization, identifying infection-derived sEVs as structurally organized immunogens rather than passive antigen reservoirs.

**Graphical Abstract:** 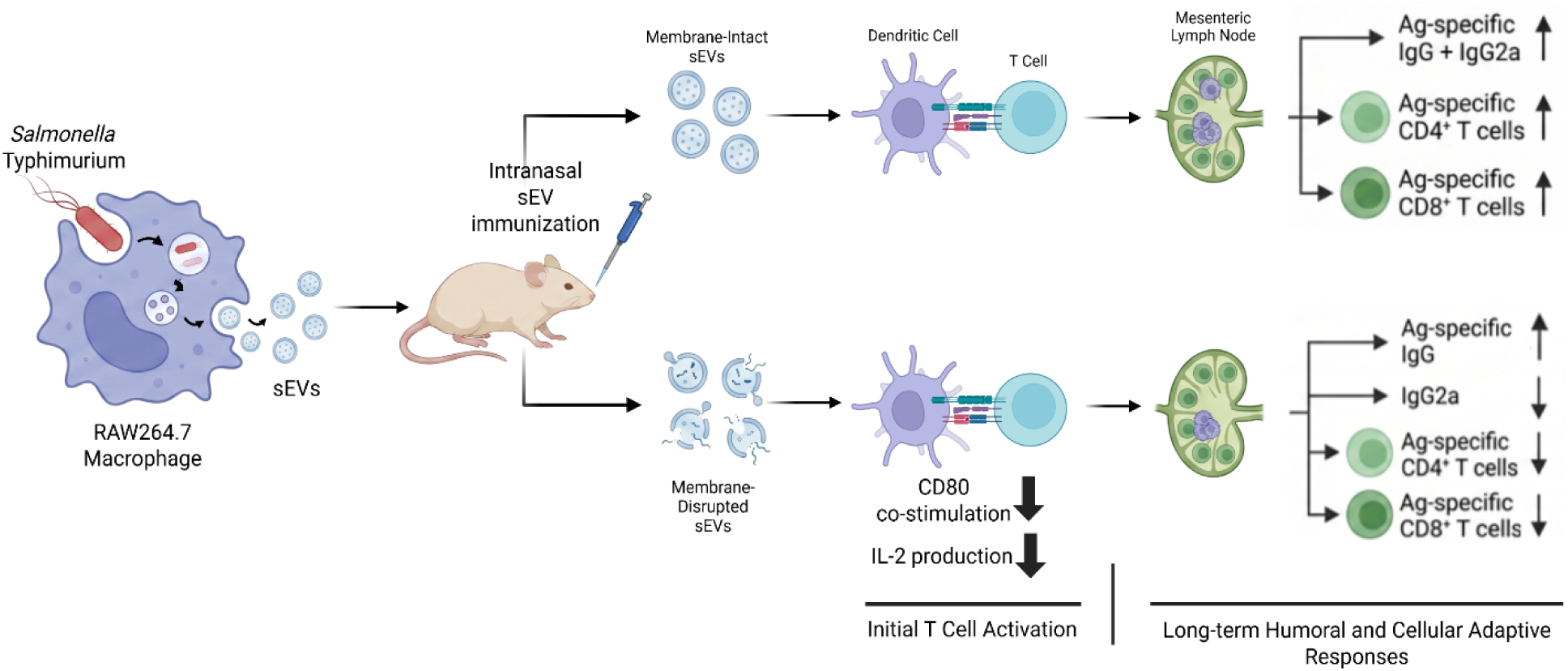

## INTRODUCTION

Non-typhoidal *Salmonella* (NTS) infections remain a major global cause of enteric and invasive disease, yet no vaccines are currently licensed for human use against NTS. Licensed *Salmonella* vaccines target typhoid fever caused by *Salmonella enterica* serovar Typhi, and are unlikely to provide broad protection against clinically important NTS serovars, including *S.* Typhimurium, because of differences in antigenic composition and immune recognition (*1,2*). Several NTS vaccine candidates are under preclinical or clinical development, including bacterial outer membrane vesicle (OMV)-based approaches, termed Generalized Modules for Membrane Antigens (GMMA), and trivalent *Salmonella* conjugate vaccines (TSCV), highlighting continued interest in multivalent strategies capable of eliciting broad immunity (*3,4*).

Protective immunity against *Salmonella* involves mucosal, humoral, and T cell–mediated responses, including antigen-specific memory capable of rapid recall following reinfection (*5*). Adoptive transfer studies have shown that serum and cellular immune components can each support protection against *Salmonella* challenge, emphasizing the need for vaccine strategies that engage multiple arms of adaptive immunity (*6,7*). Chronic or persistent *Salmonella* carriage further emphasizes the importance of durable immunological memory capable of recognizing bacterial antigens presented by antigen-presenting cells (APCs) (*8*). CD4 T cells have a particularly important role during *Salmonella* infection, where protective cellular immunity is commonly associated with T helper 1 (Th1)-type responses. Memory T cells can be broadly divided into circulating and tissue-localized populations. Circulating memory subsets include effector memory T cells (T_EM_), which can rapidly acquire effector function and traffic through lymphoid and non-lymphoid tissues, and central memory T cells (T_CM_), which preferentially home to secondary lymphoid tissues and support proliferative recall responses. Tissue-resident memory T cells (T_RM_) remain positioned at barrier sites and contribute to local protection against intracellular bacterial infections like *Salmonella* (*9,10*). These observations argue that vaccine efficacy depends not simply on antibody magnitude, but on the coordinated quality and durability of adaptive immunity.

Extracellular vesicles (EVs) have emerged as intercellular carriers of immunologically active cargo and as candidate platforms for antigen delivery. Small extracellular vesicles (sEVs), typically 30–150 nm in diameter, are released by most cell types and are present in biological fluids. They are commonly characterized by surface tetraspanins such as CD63, CD9, and CD81, as well as endosomal or cytosolic markers including Alix and TSG101 (*11,12*). During infection, sEVs may therefore provide a mechanism for packaging pathogen-derived antigens together with host molecules that influence their uptake, presentation, and immunological context. EV-associated antigens can engage adaptive immunity through several routes, including delivery of antigenic cargo to recipient APCs, transfer of peptide–MHC complexes between cells, and, under some conditions, direct presentation of vesicle-associated peptide–MHC complexes (*11,13*). Consistent with this immunological versatility, engineered and APC-derived sEVs have been shown to modulate antigen-specific responses in infectious disease and tumor models (*14–16*). Whether the physical organization of infection-derived host sEVs is itself required for effective bacterial antigen–specific immunity, however, remains for now poorly understood.

Our previous work demonstrated that sEVs isolated from *Salmonella*-infected macrophages induce long-lasting protective immunity following mucosal administration, associated with splenic memory CD4 T-cell responses (*17,18*). Proteomic analyses further identified multiple *Salmonella* antigens enriched within these vesicles, including proteins with known or predicted roles in immune recognition and protection. Individual *Salmonella* antigens such as FliC, SseB, OmpA, CirA, and SopB have been implicated in antigen-specific immune responses or protective immunity (*19–21*). These findings raised a central mechanistic question: does the intact vesicular architecture of sEVs simply transport antigen, or does it actively organize bacterial antigens and host immunoregulatory signals in a manner that determines the quality of adaptive immunity?

Here, we asked whether the structural integrity of infection-derived host sEVs governs adaptive immune programming after mucosal immunization. By comparing intact sEVs with sonication-disrupted sEVs, we show that vesicle architecture is dispensable for overall antibody induction but is selectively required for Th1-associated IgG2a responses, dendritic-cell activation and early T-cell cytokine induction, antigen-specific CD4 and CD8 memory T-cell recall, and prolonged survival after lethal *Salmonella* challenge. Together, these findings provide the first evidence that infection-derived sEVs are organized immunogenic structures whose membrane integrity coordinates humoral and cellular memory against *Salmonella*.

## RESULTS

### *Salmonella*-infected macrophages release sEVs whose membrane organization and protein compartmentalization are disrupted by sonication

sEVs released from uninfected and *Salmonella*-infected RAW264.7 macrophages were characterized using MISEV guidelines (22) to establish a model in which vesicle architecture could be perturbed without substantially altering particle size. Transmission electron microscopy confirmed the expected vesicular morphology of intact sEV preparations isolated from uninfected macrophages [sEV(−)] and *Salmonella*-infected macrophages [sEV(+)]. By contrast, sonicated vesicle [dis sEV(+)] preparations exhibited marked structural disruption and loss of intact vesicular morphology (**Fig. 1A**). Despite these morphological changes, nanoparticle tracking analysis showed similar mean particle sizes across intact and disrupted preparations (**Fig. 1B**), indicating that sonication altered vesicle architecture without substantially changing the size distribution of NTA-detectable particles.

**Figure 1:**
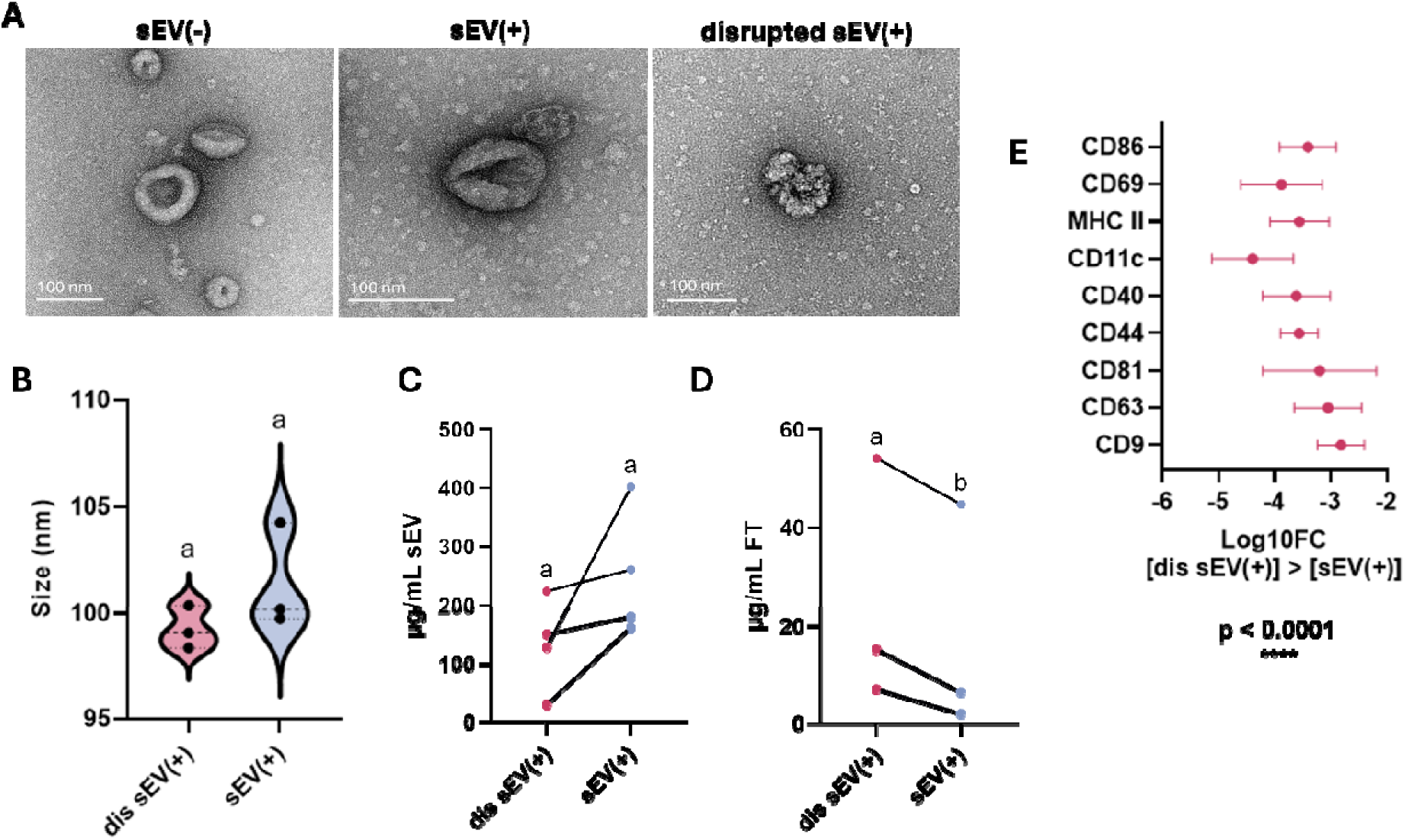
*Salmonella*-infected macrophages release nanoscale sEVs, and sonication alters vesicle-associated protein retention. **(A)** Transmission electron microscopy (TEM) images showing sEVs from uninfected macrophages [sEV(-)], intact sEVs from infected macrophages [sEV(+)], and disrupted sEVs from infected macrophages [dis sEV(+)]. **(B)** sEVs isolated from *Salmonella-*infected macrophages were either left intact or disrupted using a probe sonicator and analyzed using the ZetaView nanoparticle tracker to determine differences in particle size (nm) (n=3 independent experiments). **(C-D)** BCA assay was conducted to demonstrate whether sonication led to detachment of certain vesicle-associated protein cargo by passing the intact or disrupted sEV preparations through a 100kDa MWCO filter. Protein concentration in μg/mL is shown in retained sEV fraction **(C)** and flow through **(D)** (n=3-4 independent sEV isolations). Paired t-test analysis was conducted to determine statistical significance. **(E)** Log10 fold change in vesicle surface-associated proteins comparing dis sEV(+) versus sEV(+), determined using MACSplex mouse EV IO kit (Miltenyi) (n=4 independent experiments). Two-way ANOVA with multiple comparisons was used to determine statistical significance. All dis sEV(+) samples displayed p<0.0001 **** as compared to sEV(+) samples.

We next determined whether structural disruption altered the compartmentalization of vesicle-associated proteins. Following molecular-weight cutoff filtration, intact sEV(+) preparations contained more protein in the retained fraction, whereas sonicated sEV(+) preparations contained more protein in the flowthrough (**Fig. 1C,D**). These findings are consistent with sonication releasing proteins that were retained within or associated with intact vesicles.

The effect of sonication on the detection of vesicle-associated surface epitopes was also assessed using the MACSPlex EV assay. Intact sEV(+) preparations showed higher signals for multiple EV-associated markers than sonication-disrupted sEV(+) preparations, including CD69, MHC-II, CD11c, CD40, CD44, CD81, CD63, and CD9 (**Fig. 1E**). Because MACSPlex signal reflects both epitope accessibility and marker co-detection on captured vesicles, the reduced signal after sonication is consistent with disruption of vesicle surface organization rather than necessarily loss of individual proteins. Together, these findings establish a structural perturbation model in which sonication disrupts vesicle morphology, protein compartmentalization, and surface-marker organization while preserving the mean size of detectable particles. We therefore used intact and disrupted sEV(+) preparations to determine whether vesicle architecture influences the quality of adaptive immunity after mucosal immunization.

### Intact sEVs enhance dendritic cell costimulation and early CD4 T cell activation

To understand whether vesicle integrity influenced early dendritic cell activation and subsequent CD4 T cell responses, bone marrow-derived dendritic cells (BMDCs) were stimulated for 24 hours with 2 μg intact sEV(+) or dis sEV(+) isolated from three separate experiments (**Fig. 2A**). Intact sEV(+)-stimulation resulted in upregulation of co-stimulatory molecule CD80 (B7.1) as compared to disrupted sEV(+) stimulation. By contrast, CD86 (B7.2) and MHC-II expression did not differ significantly between intact and disrupted sEV(+)-stimulated BMDCs, although both vesicle preparations increased expression of these markers relative to unstimulated cells (**Fig. 2B-D**). Thus, disruption of vesicle architecture selectively attenuated CD80 upregulation while preserving other features of BMDC activation.

**Figure 2:**
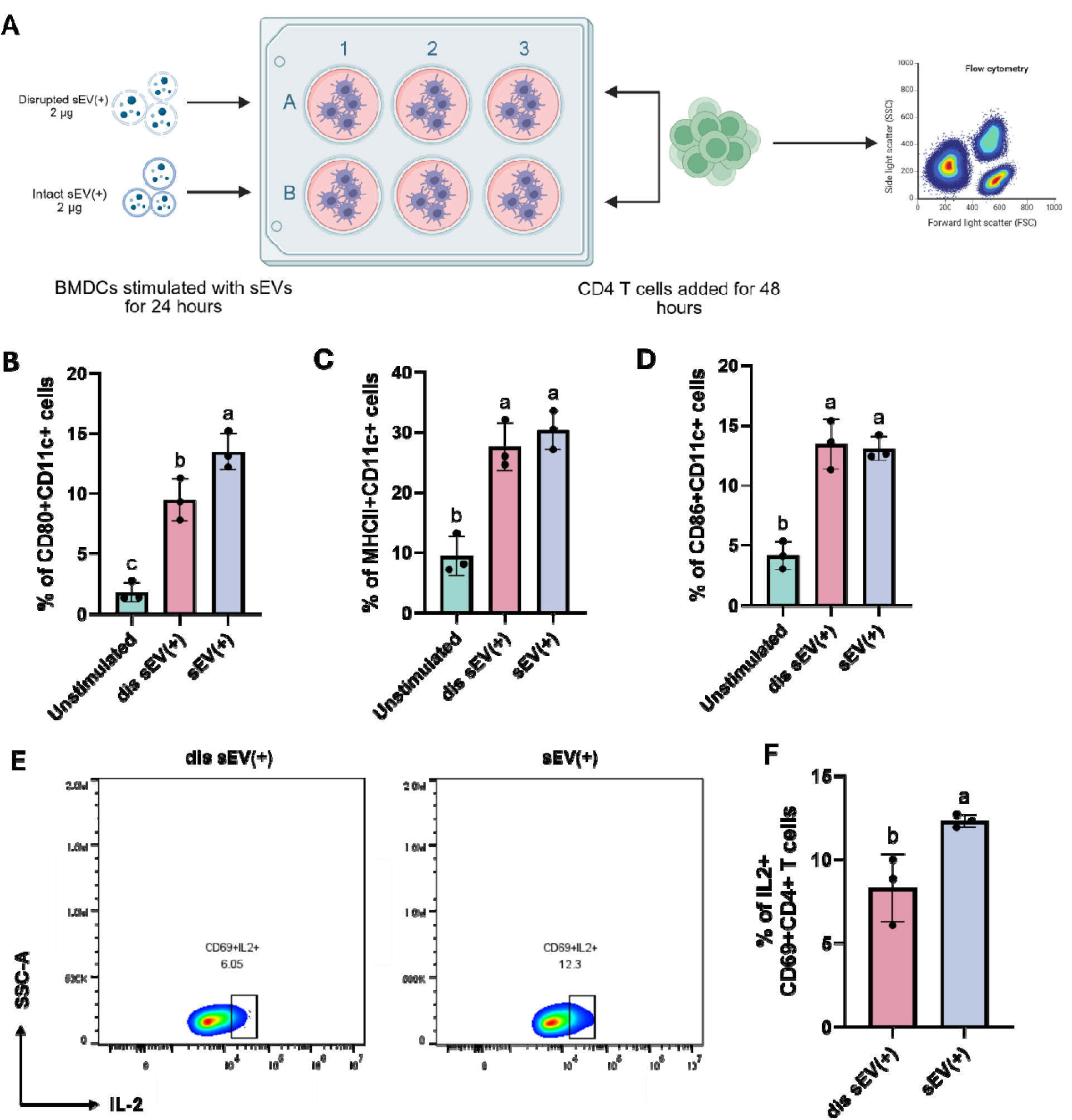
Diminished BMDC co-stimulatory signaling changes early CD4 T cell activation. **(A)** Schematic diagram showing bone marrow-derived dendritic cells (BMDCs) cultured in a 6-well plate, stimulated with 2 μg dis sEV(+) or intact sEV(+) for 24 hours, after which 3 wells containing each treatment were used for flow cytometry analysis, whereas the remaining 3 wells per treatment were used for a co-culture for 48 hours with naïve CD4 T cells isolated from Balb/c mice at a 5:1 T cell to BMDC ratio, after which flow cytometry was used to analyze T cell activation. **(B-D)** Flow cytometry analysis showing percentage of CD80+, CD86+ and MHCII+ CD11c+ dendritic cells respectively, upon no stimulation, or stimulation with dis sEV(+) or intact sEV(+). **(E)** Pseudocolor plots showing IL-2 expression of CD69+CD4+ T cells co-cultured with BMDCs stimulated with dis sEV(+) or intact sEV(+), quantified to include replicates in **(F)**. Statistical significance was determined using one-way ANOVA with post-hoc Tukey’s test or unpaired t-test and denoted as compact letter display (a,b,c). Data shown as mean **±** SEM.

We next tested whether this difference in BMDC activation was associated with altered early CD4 T cell responses. Naïve CD4 T cells were isolated from mice and co-cultured for 48 hours with intact or disrupted sEV(+) stimulated BMDCs. T cells co-cultured with BMDCs stimulated with intact sEV(+) showed increased IL-2 production by CD69+CD4+ T cells as compared to those cultured with disrupted sEV(+)-stimulated BMDCs (**Fig. 2E,F**). These findings provide a mechanistic link between intact vesicle architecture, enhanced dendritic cell costimulation, and increased early CD4 T cell activation.

### Infection-derived sEVs elicit systemic antigen-specific IgG, whereas vesicle integrity preferentially supports IgG2a responses

The immunogenicity of infection-derived sEVs was next evaluated using an intranasal prime–boost immunization strategy. BALB/c mice were immunized at weeks 0, 2, and 4 with intact sEVs from *Salmonella*-infected macrophages [sEV(+)], sonication-disrupted sEV(+) preparations [dis sEV(+)], live attenuated ΔaroA *Salmonella*, or PBS as a negative control. Serum was collected longitudinally to monitor antigen-specific IgG responses through week 7 (**Fig. 3A**). Antigen-specific humoral responses were assessed against LPS-detoxified whole *Salmonella* lysate (**Fig. S1A**) and the recombinant *Salmonella* antigens OmpA, CirA, and SopB, whose purity was confirmed before serological analysis (**Fig. S1B,C**).

**Figure 3:**
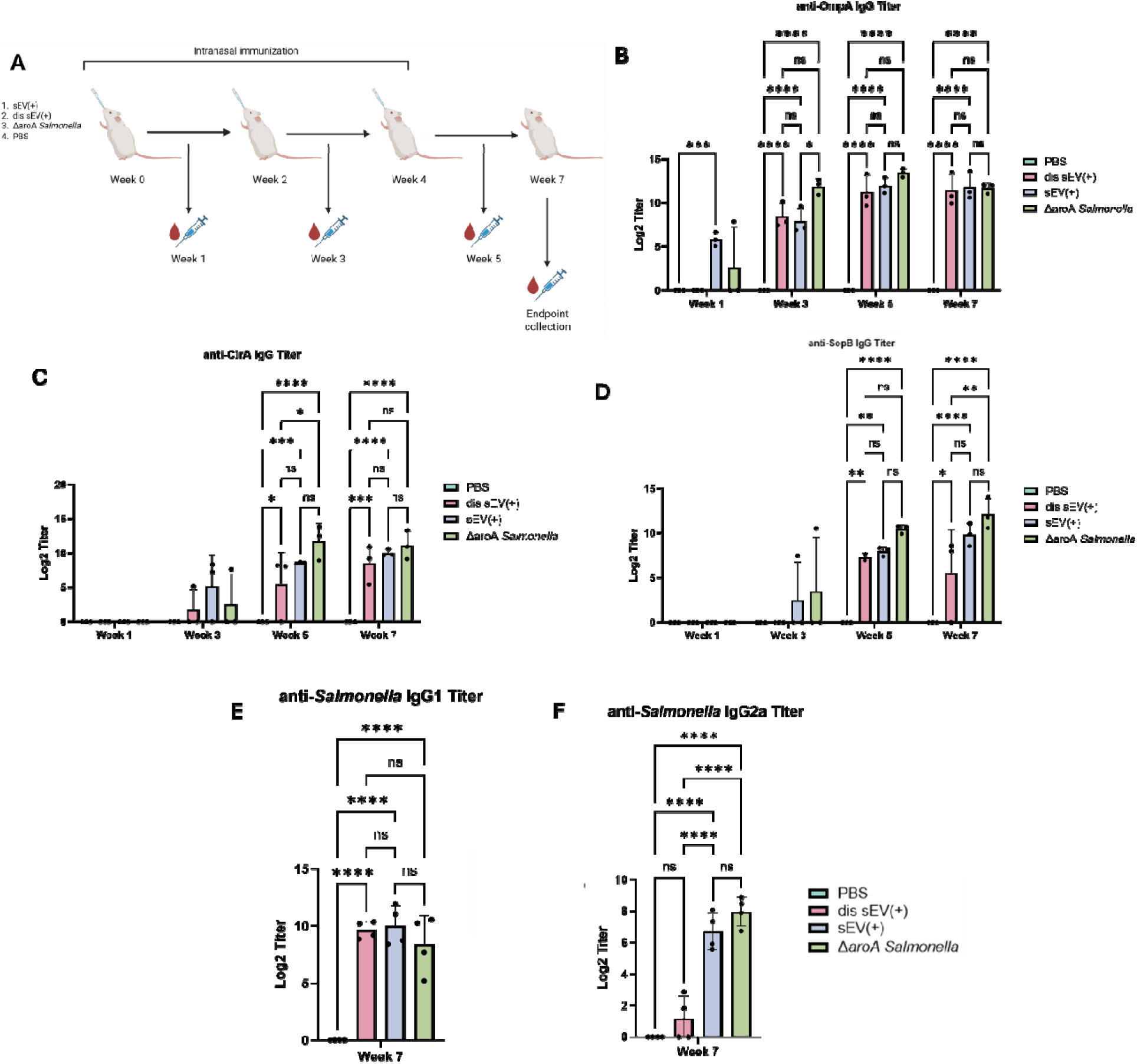
Intranasal immunization with infection-derived sEVs induces antigen-specific serum IgG responses comparable to live attenuated *Salmonella.* **(A)** Schematic diagram showing mouse immunization schedule with saphenous vein blood draws conducted at weeks 1, 3, 5, and 7 (created using Biorender). **(B-D)** Serum IgG ELISAs against *Salmonella* antigens CirA, OmpA, and SopB showing changes in Log2 titers from week 1 to week 7 post-immunization with PBS, dis sEV(+), sEV(+), or ΔaroA *Salmonella* Typhimurium UK-1 (n=3 mice per immunization group). **(E-F)** Serum IgG1 and IgG2a ELISA against LPS-detoxified whole *Salmonella* lysate showing Log2 titers 7 weeks post immunization (n=4 mice per immunization group). Two-way ANOVA test with post-hoc Tukey’s test was used to determine statistical significance. P-values have been indicated as ns (not significant), * (p<0.05), ** (p<0.01), *** (p<0.001), and **** (p<0.0001). Data shown as mean ± SEM.

Mice immunized with intact sEV(+), disrupted sEV(+), or Δ*aroA Salmonella* over time had progressively increased antigen-specific serum IgG responses relative to PBS-treated controls (**Fig. 3B–D**). Responses became evident after booster immunization and increased over time. By week 7, both intact and disrupted sEV(+) preparations elicited total antigen-specific IgG responses comparable in magnitude to those induced by live attenuated Δ*aroA Salmonella*.

We next asked whether vesicle integrity influenced IgG subclass polarization. At week 7, *Salmonella* lysate–specific IgG1 titers were comparable between mice immunized with intact and disrupted sEV(+) preparations **(Fig. 3E**). In contrast, intact sEV(+) immunization elicited significantly higher IgG2a titers than disrupted sEV(+) immunization (**Fig. 3F**). Thus, disruption of vesicle architecture did not impair overall IgG or IgG1 induction but selectively reduced the IgG2a response.

Together, these findings demonstrate that infection-derived sEVs deliver immunogenic *Salmonella* antigens sufficient to elicit robust systemic antibody responses after intranasal administration. Importantly, the sEV integrity was dispensable for the magnitude of total IgG responses but promoted IgG2a subclass polarization, suggesting that vesicle architecture influences the qualitative rather than simply quantitative features of the humoral response.

### sEV immunization enhances post-challenge mucosal IgA recognition of *Salmonella* and serum-dependent macrophage uptake

We next assessed whether antibody responses elicited by infection-derived sEVs exhibited functional activity against *Salmonella* at mucosal and systemic sites. Stool samples were collected from sEV(+)-and PBS-immunized mice following oral *Salmonella* challenge at week 7, and IgA-bound bacteria were quantified by flow cytometry. Representative pseudocolor plots showed greater IgA association with *Salmonella* in stool from sEV(+)-immunized mice than from PBS-treated controls (**Fig. 4A**). Quantification confirmed a higher frequency of IgA-bound *Salmonella* in the sEV(+)-immunized group (**Fig. 4B**). However, total *Salmonella* events recovered from stool were comparable between groups (**Fig. 4C**), indicating that the increased frequency of IgA-bound bacteria was not explained by differences in bacterial recovery.

**Figure 4:**
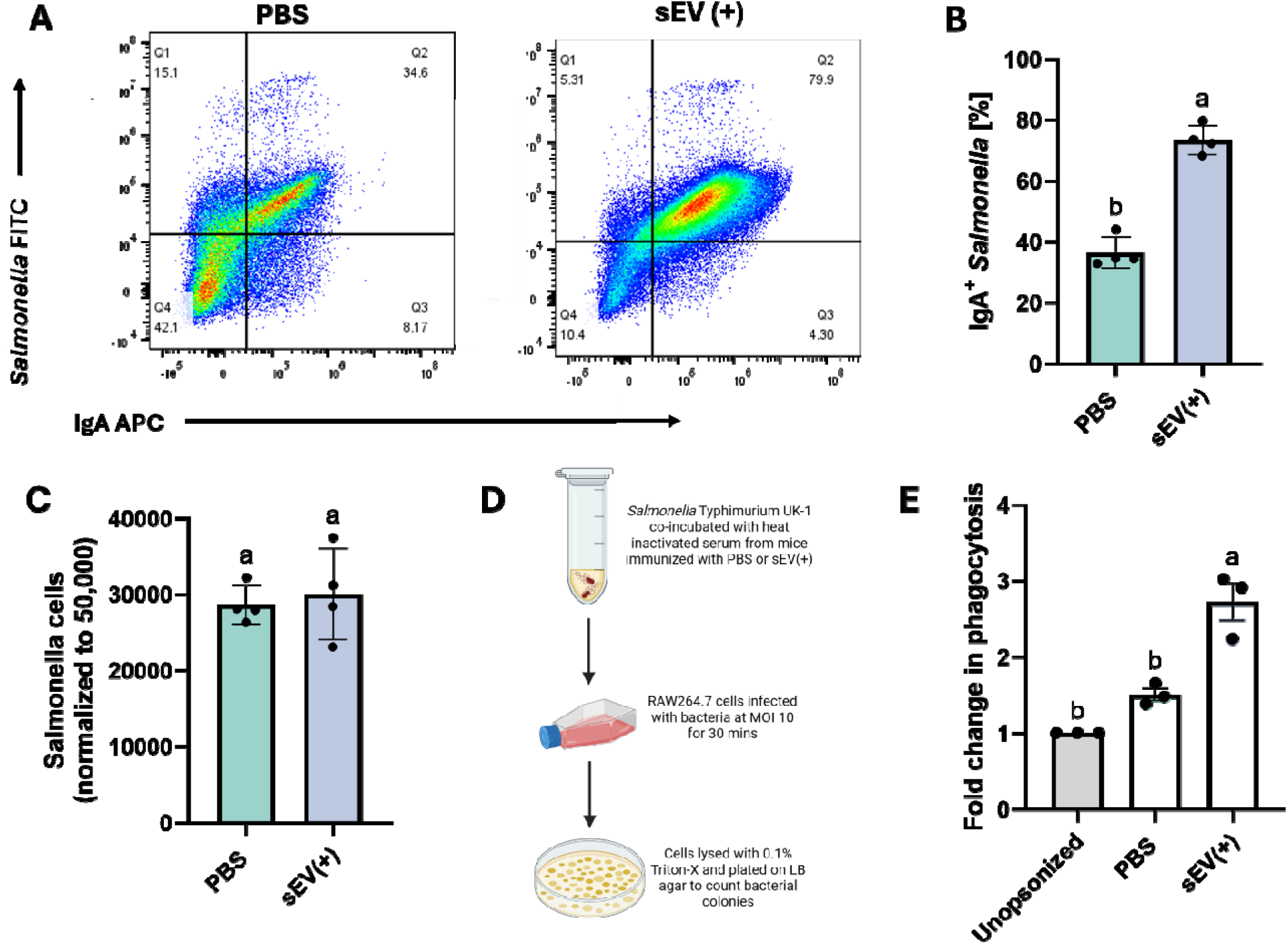
sEV immunization enhances mucosal IgA binding to *Salmonella* and promotes serum-mediated macrophage uptake. (A-B) Fecal IgA binding to *Salmonella* was demonstrated using flow cytometry **(A),** and percentage of IgA+*Salmonella* was quantified **(B)** (n=4 mice). **(C)** *Salmonella* counts in stool samples analyzed using FlowJo and normalized to 50,000 cells (n=4 mice). **(D)** Pictorial representation of opsonophagocytic assay showing bacterial co-incubation with mouse serum followed by macrophage infection and agar plating (created using Biorender). **(E)** Fold change in phagocytosis of *Salmonella* by RAW264.7 macrophages infected with either unopsonized bacteria, bacteria opsonized with serum from PBS immunized mice, or bacteria opsonized with serum from sEV(+) immunized mice (n=3 mice). One-way ANOVA test with post-hoc Tukey’s test was used to determine statistical significance. Data shown as mean **±** SEM.

We next determined whether serum elicited by sEV immunization enhanced phagocytic uptake of *Salmonella*. *Salmonella* Typhimurium UK-1 was incubated with heat-inactivated serum from PBS-or sEV(+)-immunized mice and subsequently added to RAW264.7 macrophages for 30 minutes. After gentamicin treatment to eliminate extracellular bacteria, macrophages were lysed and intracellular *Salmonella* was quantified by plating (**Fig. 4D**). Serum from sEV(+)-immunized mice significantly increased intracellular bacterial recovery relative to both un-opsonized bacteria and bacteria incubated with control serum (**Fig. 4E**), demonstrating that sEV-elicited serum components promote macrophage uptake of *Salmonella*.

Together, these findings show that intranasal sEV immunization enhances antibody-mediated recognition of *Salmonella* in the intestinal lumen after challenge and promotes serum-dependent bacterial uptake by macrophages. Thus, infection-derived sEVs elicit antibody responses with functional activity beyond increased circulating antibody titers.

### sEV immunization remodels mesenteric lymph node immune states after lethal *Salmonella* challenge

To define how sEV immunization alters immune responses in the draining lymphoid tissue after *Salmonella* challenge, sEV(+)-or PBS-immunized mice were orally challenged 7 weeks after immunization, and mesenteric lymph nodes were collected 24 hours later for single-cell RNA sequencing (scRNA-seq). Integrated analysis of mesenteric lymph node cells by scRNA-seq identified multiple lymphoid and antigen-presenting cell populations, including naïve CD4 T cells, central memory CD4 T cells (T_CM_), cytotoxic CD8 T cells, mature B cells, activated/effector memory CD8 T cells, regulatory T cells, antigen-experienced B cells, proliferating cells, and conventional dendritic cells (cDCs) (**Fig. 5A, S2A**). The distribution of transcriptionally defined immune populations differed between groups after challenge, with greater representation of central memory CD4 T cells, effector CD8 T cells, and mature B cells in sEV(+)-immunized mice (**Fig. 5A, S2A**).

**Figure 5:**
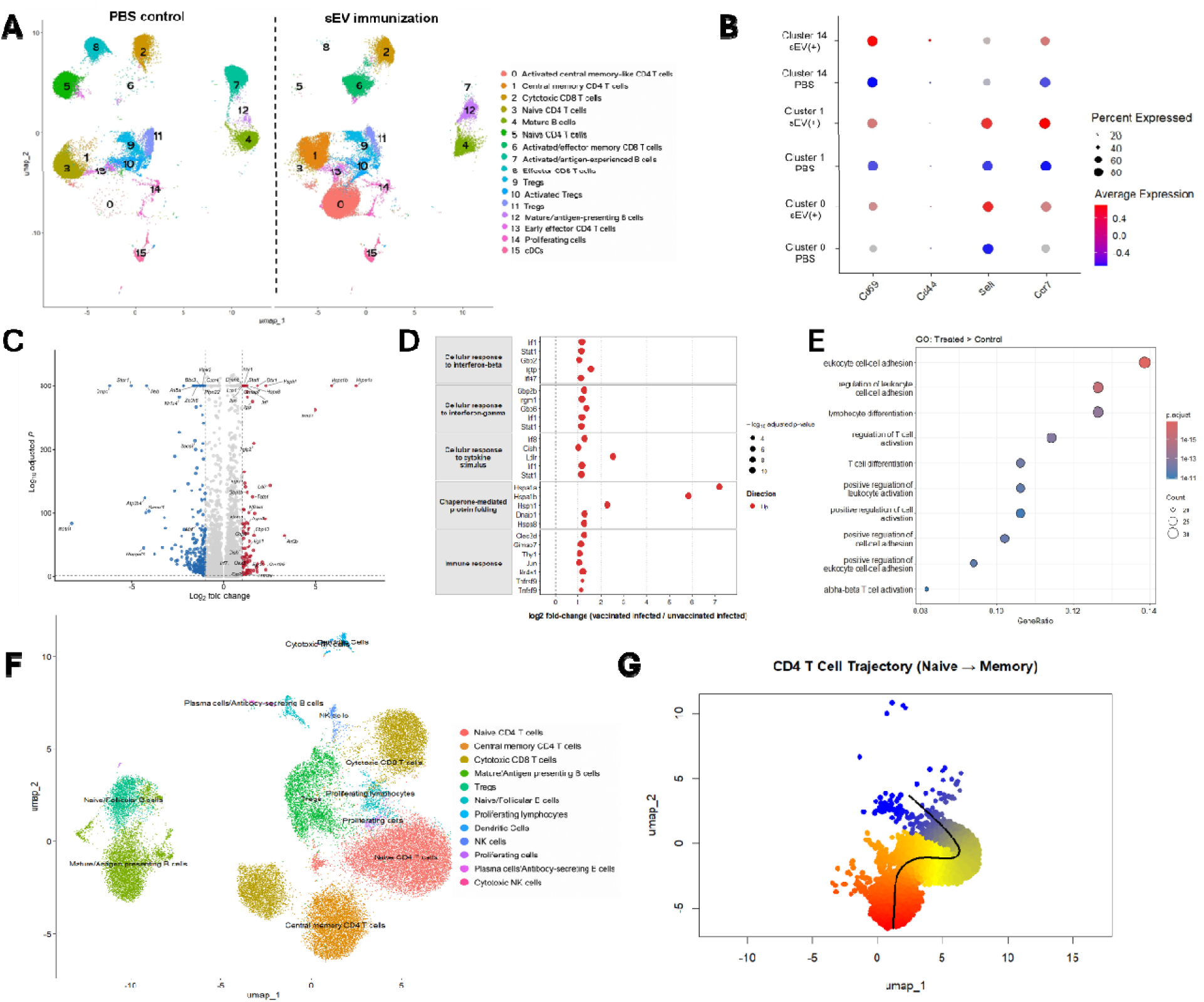
sEV immunization remodels mesenteric lymph node immune states after lethal *Salmonella* challenge. **(A)** Uniform Manifold Approximation and Projection (UMAP) plots of mesenteric lymph node (mLN) cells from PBS-and sEV(+)-immunized mice 24 hours after lethal *Salmonella* challenge at week 7 post-immunization (n=2 mice per condition). **(B)** Percent expression of Cd69, Cd44, Ccr7, and Sell (Cd62L) in all CD4 T cell clusters (excluding Tregs) in both PBS-and sEV(+)-immunized mice. **(C)** Volcano plot showing the expression data across all CD4 T cell clusters. **(D)** Select gene expression analysis from **(C). (E)** GO enrichment pathway analysis conducted on all CD4 T cell clusters (excluding Tregs) on R studio showing activated pathways in treated [sEV(+)] versus control (PBS) samples. **(F-G)** Uniform Manifold Approximation and Projection (UMAP) plot showing clustering of different cell populations in murine mesenteric lymph nodes (mLNs) of sEV(+)-immunized mice **(F)**, and Slingshot trajectory prediction based on UMAP plot clusters in **(F)** from naïve to memory (blue to red) CD4 T cells **(G)**.

Within the central memory CD4 T cell cluster, cells from sEV(+)-immunized mice exhibited increased expression of *Cd69*, *Cd44*, *Ccr7*, and *Sell* relative to PBS controls (**Fig. 5B, Fig. S2B,C**). The concurrent expression of activation-and antigen-experience-associated genes (*Cd69* and *Cd44*) with the lymphoid-homing genes *Ccr7* and *Sell* was consistent with an activated central memory–associated transcriptional state following challenge.

Differential expression analysis across all CD4 T cell clusters, excluding regulatory T cells (Tregs), showed broad transcriptional differences associated with prior sEV immunization (**Fig. 5C; Table S1**). CD4 T cells from sEV(+)-immunized mice exhibited increased expression of TCR-responsive and activation-associated genes, including *Nr4a1, Egr2, Egr3, Dtx1, Jun, Cish, Nfkbid, Tnfrsf9, Gimap7,* and *Thy1*. These changes were consistent with enhanced antigen-responsive signaling and activation of the CD4 T cell compartment after *Salmonella* challenge.

sEV(+) immunization was also associated with increased expression of interferon-responsive and cell-intrinsic antimicrobial genes within CD4 T cell clusters, including *Stat1, Irf1, Oas3, Gbp10, Gbp6, Gbp2b, Igtp, Tgtp1, Tgtp2, Iigp1, Irgm1*, and *Irgm2*. Pathway-level analysis highlighted transcriptional modules associated with cellular responses to type I interferon, interferon-γ, cytokine stimulation, immune response, and chaperone-mediated protein folding (**Fig. 5D**). KEGG pathway mapping also supported differential regulation of cytokine signaling in CD4 T cell clusters from sEV(+)-immunized mice, with increased expression of *Stat1* across multiple cytokine-and interferon-associated JAK–STAT signaling nodes, together with increased *Cish, Myc*, and *Ccnd2* (**Fig. S2D**). Together, these analyses identified coordinated TCR-, cytokine-, and interferon-responsive transcriptional programs in CD4 T cells from sEV(+)-immunized mice following challenge.

Gene ontology analysis independently supported this phenotype. Genes enriched in non-Treg CD4 T cells from sEV(+)-immunized mice mapped to processes associated with leukocyte cell–cell adhesion, lymphocyte differentiation, regulation of T cell activation, and T cell differentiation (**Fig. 5E**). In contrast, pathways enriched in control T cell clusters were associated primarily with translation, ribosome biogenesis, and electron transport processes (**Fig. S2E**). Together, these pathway-level analyses identified a shift toward activation-, differentiation-, and immune interaction–associated transcriptional programs following sEV immunization.

We next examined relationships among CD4 T cell states using pseudotime trajectory analysis. Slingshot modeling identified an inferred trajectory from naïve CD4 T cell states toward memory-associated CD4 T cell states in sEV(+)-immunized mice (**Fig. 5F,G**). A comparable memory-associated trajectory was not evident in *Salmonella*-challenged PBS controls (**Fig. S3A,B**). These findings were consistent with preferential representation of transcriptional states associated with CD4 T cell activation and memory differentiation following sEV immunization.

B cell-focused analysis of the scRNA-seq data also supported adaptive immune remodeling after sEV(+) immunization. Genes enriched in B cell clusters from sEV(+)-immunized mice were specifically associated with regulation of immune effector processes, lymphocyte differentiation, B cell activation, leukocyte proliferation, and positive regulation of lymphocyte activation (**Fig. S4A; Table S2**). In contrast, genes enriched in PBS control B cells were associated with DNA repair, DNA recombination, macroautophagy, and protein polyubiquitination pathways unrelated to immune cell activation (**Fig. S4B**). Differential expression analysis identified increased expression of interferon-and antibacterial response–associated genes, including *Irf1*, *Stat1*, *Gbp2*, *Tlr3*, *Gbp7*, *Irgm1*, *Isg15*, *Nod2*, *Ly6a*, and *Gbp10*, in B cells from sEV(+)-immunized mice (**Fig. S4C,D**). Thus, prior sEV immunization also altered B cell transcriptional programs during the early response to *Salmonella* challenge.

Together, these data show that sEV immunization reshapes the early mesenteric lymph node response to *Salmonella*, with increased representation of memory-associated lymphocyte states and enhanced activation-, TCR-, and interferon-responsive programs in CD4 T and B cell populations. These findings imply that infection-derived sEV immunization establishes distinct adaptive immune states that are rapidly engaged upon pathogen re-exposure.

### Intact sEV immunization primes antigen-specific CD4 and CD8 memory T cell recall in mesenteric lymph nodes

We next determined whether vesicle integrity influenced functional T cell memory against defined *Salmonella* antigens carried by infection-derived sEVs. BALB/cJ mice were intranasally immunized at weeks 0, 2, and 4 with intact sEV(+), disrupted sEV(+) [dis sEV(+)], or PBS. At week 7, mice were orally challenged with lethal *Salmonella*, and mesenteric lymph nodes were collected 24 hours later. Single-cell suspensions were stimulated *ex vivo* for 72 hours with a recombinant antigen cocktail containing OmpA, SopB, CirA, FliC, and OmpD, all previously identified in sEVs from *Salmonella*-infected macrophages (*18*), followed by intracellular cytokine analysis (**Fig. S5A**).

Antigen restimulation elicited robust cytokine recall responses in memory CD4 and CD8 T cells from mice immunized with intact sEV(+). Representative flow cytometry plots, based on our indicated gating strategy (**Fig. S5B**), showed increased frequencies of IFNγ+ and IL-2+ CD44+CD62L+ CD4 T_CM_ cells following antigen-cocktail restimulation in cells from intact sEV(+)-immunized mice (**Fig. 6A,D**). Quantification confirmed increased percentages and numbers of IFNγ+ and IL-2+ CD4 T_CM_ cells in the intact sEV(+)-immunized group (**Fig. 6B,C,E,F**). CD44+CD62L− effector memory (T_EM_) CD4 T cells from intact sEV(+)-immunized mice also showed increased IFNγ and IL-2 production after antigen-cocktail stimulation (**Fig. S6A,B**). In contrast, disrupted sEV(+) immunization did not induce comparable antigen-responsive IFNγ+ or IL-2+ CD4 T_CM_ and IFNγ+ or IL-2+ CD4 T_EM_ cell responses, indicating that vesicle integrity is important for generating functional CD4 T cell recall responses to sEV-associated *Salmonella* antigens (**Fig. 6A-F; Fig. S7A,B**). Thus, preservation of vesicle architecture favored the development of cytokine-competent, antigen-responsive CD4 memory T cells.

**Figure 6:**
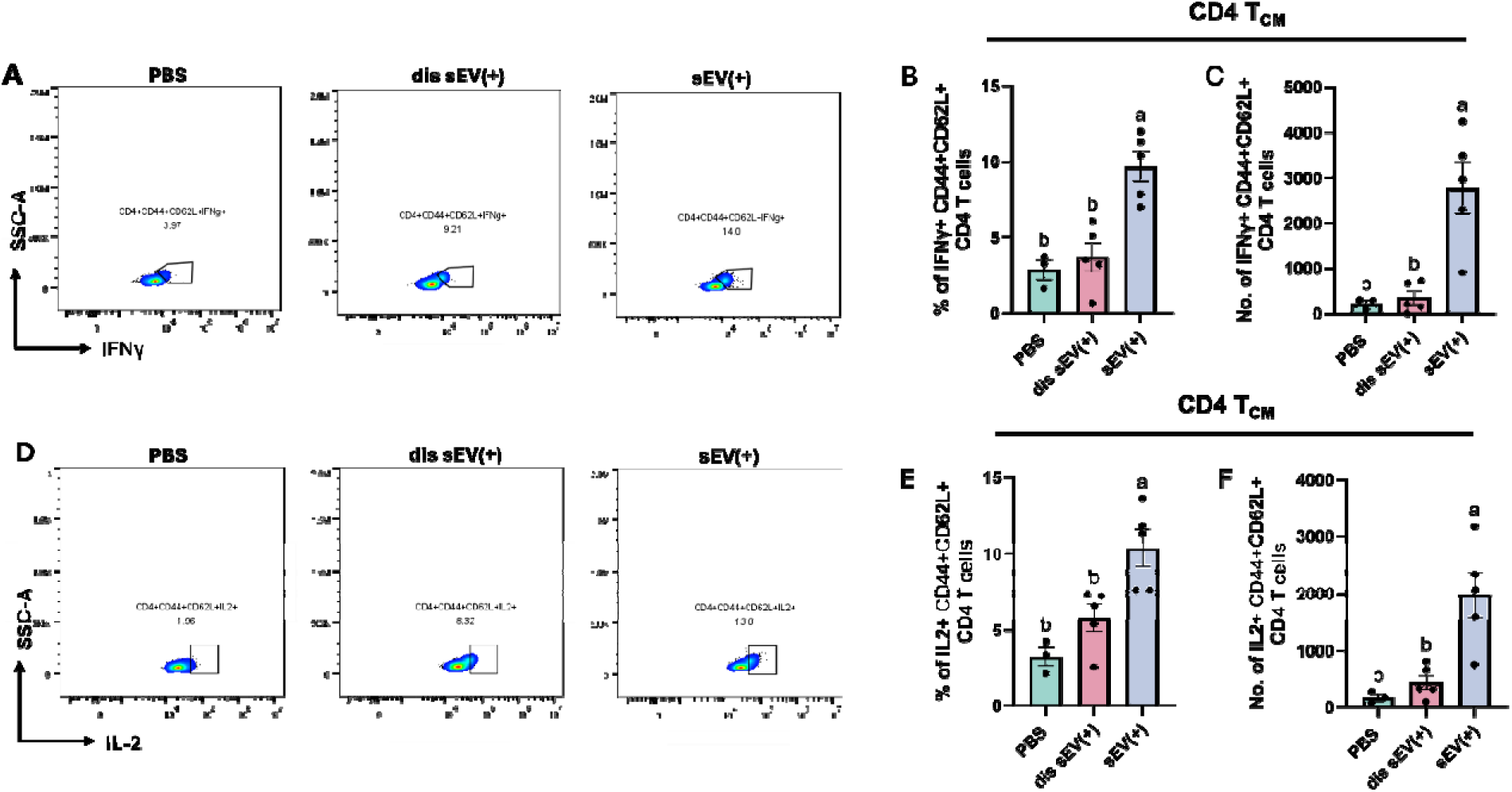
Mesenteric lymph nodes of mice immunized with sEVs activate *Salmonella* antigen-specific central memory (T_CM_) CD4 T cells. **(A)** Representative flow cytometry pseudocolor plots showing percentage of IFNγ+ central memory (T_CM_) CD4 T cells (CD4+CD44+CD62L+). **(B)** Percentage of IFNγ-expressing CD44+CD62L+ (T_CM_) CD4 T cells. **(C)** Number of IFNγ-expressing CD44+CD62L+ (T_CM_) CD4 T cells. **(D)** Representative flow cytometry pseudocolor plots showing percentage of IL2+ central memory (T_CM_) CD4 T cells (CD4+CD44+CD62L+). **(E)** Percentage of IL2-expressing CD44+CD62L+ (T_CM_) CD8 T cells. **(F)** Number of IL2-expressing CD44+CD62L+ (T_CM_) CD4 T cells. (n=3-5 mice per immunization group). Statistical significance was determined using one-way ANOVA with post-hoc Tukey’s test, shown as compact letter display (a,b).

A similar dependence on vesicle integrity was observed in the CD8 T cell compartment. Restimulation with the defined *Salmonella* antigen cocktail increased IFN-γ–producing CD8 T cells from intact sEV(+)-immunized mice relative to PBS controls and was accompanied by increased IL-2 production. These responses were substantially reduced or absent following immunization with disrupted sEV(+) (**Fig. 7A–F**). These data demonstrate that intact infection-derived sEVs prime recall responses to defined vesicle-associated *Salmonella* antigens in both CD4 and CD8 T cell compartments.

**Figure 7:**
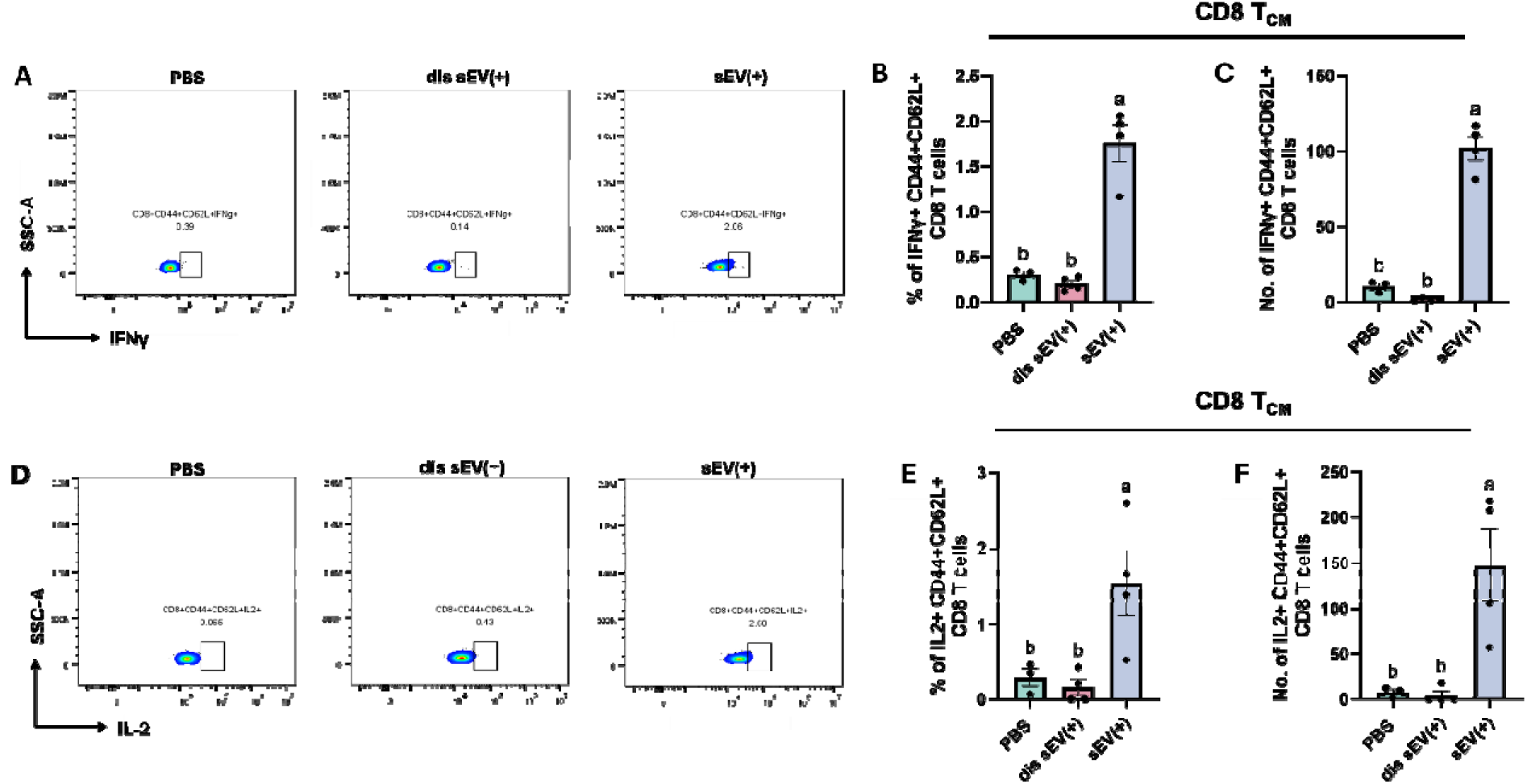
Mesenteric lymph nodes of mice immunized with sEVs activate *Salmonella* antigen-specific central memory (T_CM_) CD8 T cells. **(A)** Representative flow cytometry pseudocolor plots showing percentage of IFNγ+ central memory (T_CM_) CD8 T cells (CD8+CD44+CD62L+). **(B)** Percentage of IFNγ-expressing CD44+CD62L+ (T_CM_) CD8 T cells. **(C)** Number of IFNγ-expressing CD44+CD62L+ (T_CM_) CD8 T cells. **(D)** Representative flow cytometry pseudocolor plots showing percentage of IL2+ central memory (T_CM_) CD8 T cells (CD8+CD44+CD62L+). **(E)** Percentage of IL2-expressing CD44+CD62L+ (T_CM_) CD8 T cells. **(F)** Number of IL2-expressing CD44+CD62L+ (T_CM_) CD8 T cells. (n=3-5 mice per immunization group). Statistical significance was determined using one-way ANOVA with post-hoc Tukey’s test, shown as compact letter display (a,b).

We next examined whether vesicle integrity was associated with altered antigen-presenting cell activation in vivo. MHC-II expression on dendritic cells was comparable among immunization groups, whereas CD80 and CD86 expression was increased on mesenteric lymph node dendritic cells from mice immunized with intact sEV(+) relative to disrupted sEV(+)-and PBS-immunized mice (**Fig. S6C-E**). These data link intact sEV immunization with enhanced dendritic cell costimulatory capacity, consistent with the stronger antigen-specific T cell recall responses observed in the same draining lymphoid compartment.

To assess the anatomical distribution of antigen-responsive recall, we performed parallel analyses in axillary lymph nodes. Restimulation with the defined *Salmonella* antigen cocktail did not elicit significant IFN-γ or IL-2 responses in axillary lymph node T cells, nor increased the number of central memory CD4 T cells (**Fig. S6F-H**), suggesting that the antigen-responsive memory T cell activation observed after sEV immunization was most prominent in mesenteric lymph nodes.

We independently assessed recall responses to vesicle-associated antigenic material by restimulating mesenteric lymph node cells with intact sEV(+), disrupted sEV(+), or sEVs from uninfected macrophages [sEV(−)] for 72 hours (**Fig. S7A**). In lymphoid cells from sEV(+)-immunized mice, *ex vivo* stimulation with intact or disrupted sEV(+) preparations induced a higher frequency of IFNγ+ and IL-2+ CD4 T_CM_ cells compared with stimulation using sEV(−) preparations or unstimulated controls (**Fig. S7B,C**). A similar increase was observed for IFNγ production by CD4 T_EM_ cells (**Fig. S7D**). When compared to mice immunized with PBS, *ex vivo* stimulation with intact sEV(+), disrupted sEV(+), and sEVs from uninfected macrophages [sEV(-)] did not lead to significant IFNγ or IL-2 production by either T_CM_ or T_EM_ cells (**Fig. S7B-D**). Thus, vesicle restimulation alone provided limited evidence for immunization-dependent recall, whereas the defined *Salmonella* antigen cocktail revealed a clear antigen-specific memory response.

Together, these findings show that intact sEV immunization primes antigen-specific CD4 and CD8 memory T cells capable of rapid IFN-γ and IL-2 recall after *Salmonella* challenge. Disruption of vesicle architecture markedly impaired these responses, linking vesicle integrity to the development of functional cellular memory and enhanced dendritic cell costimulatory activation.

### Intact sEV(+) immunization prolongs survival following lethal *Salmonella* challenge

We finally determined whether the immune responses elicited by intact sEVs translated into improved protection against lethal infection. Immunized mice were challenged with *Salmonella* (4.5 x 10^6^ CFUs) and monitored through the experimental endpoint. Immunization with intact sEV(+) prolonged survival compared with disrupted sEV(+)-and PBS-immunization (**Fig. 8A**). Clinical condition scores revealed a corresponding benefit, with intact sEV(+)-immunized mice showing reduced clinical deterioration compared with PBS-or disrupted sEV(+)-immunized mice (**Fig. 8B**).

**Figure 8:**
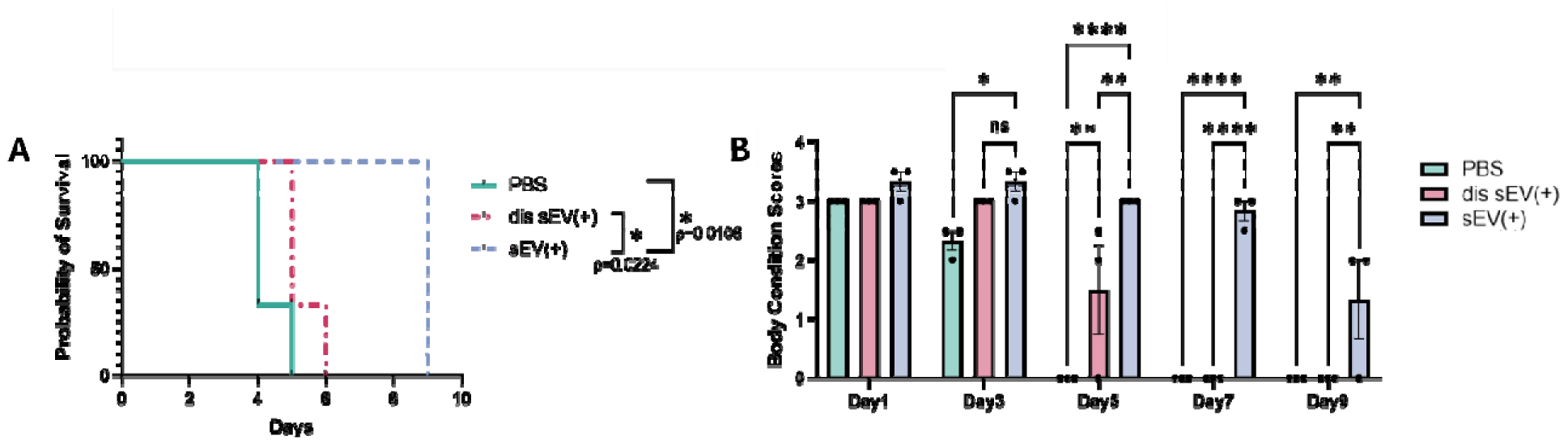
Intact sEV(+) immunization prolongs survival following lethal *Salmonella* challenge. **(A)** Kaplan-Meier survival curve showing survival of mice orally challenged with a lethal dose of *Salmonella* Typhimurium UK-1 (4.5 x 10^6^ CFUs) at week 7, after three rounds of immunization with either PBS, disrupted sEV(+), or intact sEV(+). Log rank Mantel-Cox test was used to determine statistical significance. (n=3) **(B)** Body condition scores (BCS) of mice lethally challenged with *Salmonella* 7 weeks after immunization with either PBS, dis sEV(+), or intact sEV(+), measured over a span of 9 days following a BCS scale from 1 to 5. (n=3).

Together, these findings demonstrate that preservation of sEV architecture is associated with enhanced protection against lethal *Salmonella* challenge. When considered with reduced dendritic cell costimulation observed after vesicle disruption, the impaired IgG2a response, and diminished antigen-specific T cell recall, these data identify intact vesicle organization as an important determinant of the quality and protective capacity of sEV-induced immunity.

## DISCUSSION

Here, we identify infection-derived host sEVs as structured antigen carriers that coordinate humoral, mucosal, and memory T cell immunity after intranasal immunization. Using *Salmonella* infection as a model of enteric bacterial immunity, we show that sEVs produced by infected macrophages elicit antigen-specific serum IgG, enhance mucosal IgA binding to *Salmonella* after challenge, promote serum-dependent macrophage uptake, and remodel immune states in mesenteric lymph nodes after lethal infection. Intact sEVs also enhanced BMDC CD80 expression and subsequent IL-2 production by cocultured CD69^+^ CD4 T cells, linking vesicle integrity to dendritic cell co-stimulation and early T cell activation. Most importantly, we show that disrupted sEVs fail to generate comparable IgG2a production and antigen-specific CD4 and CD8 memory T cell recall. Thus, infection-derived sEVs do not function merely as antigen reservoirs. Rather, their intact architecture appears to preserve antigen organization and APC-instructive surface features that support cellular immune programming.

Previous work from our group demonstrated that sEVs isolated from *Salmonella*-infected macrophages enhance protection against lethal *Salmonella* challenge and induce long-term immunity associated with pathogen-specific secretory IgA, serum IgG, and memory CD4 T cell responses in splenic tissues (*17,18*). However, whether intact vesicle architecture was required for these effects, how sEV immunization shaped memory-associated immune states in draining lymphoid tissues, and whether cellular recall responses were directed against defined antigens carried by infection-derived sEVs remained unresolved. The present study addresses these questions by comparing intact and sonication-disrupted sEVs and by testing recall responses to a defined set of sEV-associated *Salmonella* antigens, including OmpA, SopB, CirA, FliC, and OmpD.

Sonication did not eliminate nanoscale particles, but it altered sEV morphology, redistributed vesicle-associated protein cargo, and reduced detection of multiple EV-associated surface markers. These findings indicate that disrupted sEV preparations retain antigenic material but lose key features of vesicle organization, membrane-associated protein display, and surface architecture that may be required for efficient cellular immune priming. Importantly, filtration was used to assess protein redistribution after sonication but was not applied to preparations used for immunization, maintaining equivalent administered total sEV protein doses between intact and disrupted sEV groups. This distinction was mirrored functionally: both intact and disrupted sEV(+) preparations elicited antigen-specific serum IgG, whereas analysis of IgG subclasses revealed significantly diminished IgG2a responses after disrupted sEV(+) immunization. In mice, IgG2a is commonly associated with Th1-skewed immune responses, whereas IgG1 is more often associated with Th2-type cytokine environments (23). Oral vaccination using live attenuated *Salmonella* has been shown to elicit increased IgG2a production (24). Furthermore, only intact sEV(+) immunization generated robust antigen-directed memory T cell recall. Thus, vesicle-associated antigenic cargo remained sufficient for total IgG induction, whereas intact vesicle architecture preferentially supported IgG2a responses and cellular recall.

The vesicle-associated markers showing reduced detection after sonication provide a plausible mechanistic link between vesicle architecture and memory T cell priming. Reduced detection of tetraspanins CD9, CD63, and CD81 is consistent with altered vesicle membrane organization and may influence interactions with recipient APCs. In parallel, reduced detection of immune-associated proteins such as MHC II and CD40 suggests altered organization or accessibility of molecules involved in antigen presentation and costimulatory signaling. MACSPlex detection depends on epitope accessibility and marker co-detection, however, reduced signal cannot distinguish physical loss of individual proteins from altered vesicle surface organization. Previous studies have shown that CD4 T cell interactions with EVs derived from CD40/IL-4–stimulated chronic lymphocytic leukemia B cells can enhance proliferation, migration, and immune synapse signaling (25). EVs bearing agonistic CD40 antibodies have been shown to induce BMDC activation and release of IL12-p70 more efficiently than unconjugated CD40 antibody (26) whereas CD11c, CD44, and CD69 may further contribute to APC targeting, adhesion, activation state, or intercellular communication (27,28). Our BMDC experiments provide additional functional support for a relationship between vesicle architecture and APC activation. Intact sEVs induced greater CD80 expression than disrupted sEVs, despite comparable MHC-II and CD86 expression between the two vesicle-treated groups, and BMDCs exposed to intact sEVs subsequently elicited greater IL-2 production by cocultured CD69^+^ CD4 T cells. These findings link vesicle integrity with enhanced dendritic cell costimulation and early CD4 T cell activation, although they do not establish CD80 as the causal mediator of this response.

The whole-vesicle restimulation experiments require a more cautious interpretation. Although intact and disrupted sEV(+) preparations retained antigenic material capable of stimulating cytokine production *ex vivo*, these responses did not significantly differ from those observed in equivalently stimulated cells from PBS-immunized mice. Thus, whole-vesicle restimulation alone does not establish immunization-dependent memory recall. In contrast, restimulation with the defined *Salmonella* antigen cocktail revealed clear antigen-specific CD4 and CD8 recall responses that were preferentially generated by intact sEV immunization. In vivo, intact sEVs may therefore deliver bacterial antigens together with host-derived molecular cues that promote efficient APC engagement and memory T cell programming. DC-derived EVs bearing antigen or peptide complexes have been demonstrated to activate naïve CD4 T cells both in vivo and in vitro. Additionally, EVs have been shown to support antigen-specific T cell activation in the presence of mature DCs lacking MHC II (29).

In this model, disrupted sEVs preserve antigenic cargo sufficient for total IgG and IgG1 antibody induction, whereas IgG2a responses and antigen-specific CD4 T cell recall are markedly impaired. While activation of follicular helper T cells (T_FH_) has not been directly characterized in this study, the selective reduction in IgG2a together with the Th1-associated cellular responses observed here is consistent with altered immune polarization after vesicle disruption. The relative contributions of T_FH_ and other CD4 T cell subsets to this antibody phenotype remain to be determined.

These findings are also relevant to broader EV engineering strategies, because sonication is often used to promote cargo loading into EVs or to disperse aggregated vesicle preparations before downstream use (30,31). Although sonication can improve cargo incorporation, particle dispersion, or cellular uptake, our data indicate that it can also compromise surface epitope retention, morphology, and sEV function.

The defined antigen-cocktail recall responses observed here support the concept that infection-derived sEVs deliver bacterial antigens in an immunologically functional form. Several antigens included in the recall cocktail have been implicated in *Salmonella* immune recognition, bacterial fitness, or protective immunity (32–34). FliC is a flagellar antigen with strong immunostimulatory activity, and *Salmonella* surface or secreted antigens, including SseB and FliC, have been shown to enhance protective immunity in mouse vaccination models (32,35). OmpA can activate dendritic cells and promote a Th1-skewed response characterized by increased IFNγ and reduced IL-4 production (35). CirA, an iron-regulated outer membrane receptor, contributes to *S.* Typhimurium survival or colonization in infection models and has been reported to confer protection after immunization (33,34). SopB is a virulence-associated effector, and attenuated *Salmonella* strains carrying an additional *sopB* mutation have been associated with increased dendritic cell expression of co-stimulatory molecules, including CD80 and CD86 (32). Together, these studies support the selection of OmpA, SopB, CirA, FliC, and OmpD as a defined antigen set to test whether intact sEV immunization generates functional recall responses against biologically relevant *Salmonella* cargo. The ability of intact sEV immunization to generate IFNγ-and IL-2-producing memory T cell responses against this antigen set indicates that sEV-associated bacterial cargo can be processed and presented as functional recall targets. Importantly, this response was most evident in mesenteric lymph nodes compared with axillary lymph nodes, consistent with the relevance of gut-draining lymphoid tissues during enteric *Salmonella* challenge and with prior evidence that central memory T cells can persist in mesenteric lymph nodes after *Salmonella* exposure (36).

The preservation of cellular recall responses after intact sEV immunization may be especially relevant for intracellular bacterial pathogens, for which durable T cell immunity is often required for effective protection. CD4 T cells producing IFNγ, IL-2, and TNFα have been implicated in protective responses against intracellular pathogens, including *Leishmania major* and *Mycobacterium tuberculosis* (37,38). Furthermore, comparison of effector and central T cell population in the context of *L. major* immunity have shown that the absence of persistent pathogen can lead to loss of short-lived effector T cells while permitting preservation of long-lived T_CM_ cells (39). Although the present study did not directly assess tissue-resident memory T cells, prior work in mucosal or intracellular infection models has shown that vaccination can generate tissue-localized memory populations with rapid protective capacity. For example, skin-resident CD4 TRM cells have been characterized in murine leishmaniasis, where they provide early protection through recruitment of inflammatory monocytes and CX3CR1-dependent circulating T cells (40,41). Whether infection-derived sEVs similarly generate intestinal or mucosal T_RM_ populations after intranasal immunization remains an open question for future studies. The route of sEV delivery may also influence the quality and anatomical distribution of immunity, as previous work showed that orally administered sEVs from infected macrophages can elicit protective responses in vivo as well (42). Comparative studies of mucosal delivery routes may clarify how route shapes local antibody responses, tissue-resident memory, and systemic recall immunity.

Taken together, these findings show that sEVs released from *Salmonella*-infected macrophages act as structured mucosal immunogens that coordinate antibody responses, mesenteric lymph node remodeling, and antigen-directed memory T cell recall. The divergence between preserved humoral immunogenicity and impaired cellular recall after vesicle disruption identifies vesicle integrity as a determinant of adaptive immune quality. These findings support infection-derived host sEVs as candidate nonconventional mucosal vaccine platforms and provide a foundation for designing vesicle-based approaches that preserve antigen organization and APC-instructive signals to promote durable immunity against enteric bacterial pathogens.

### Limitations of the study

This study has several limitations that merit consideration. The antigen cocktail used here was based on selected *Salmonella* proteins previously identified in infection-derived sEVs (*18*), but it does not capture the full complexity of vesicle-associated bacterial cargo or the contribution of individual antigens. Although recombinant Salmonella antigens can elicit cytokine responses and protective immunity in some models (19,20,43–46), they may not recapitulate the antigen organization or APC-instructive signals provided by intact sEVs. In addition, the APC subsets, uptake receptors, and vesicle-associated host molecules that mediate this effect remain to be defined. Finally, although sonication is a well-described strategy to disrupt vesicle integrity (31), it is not selective for a single structural feature. The impaired cellular recall observed after sonication may therefore reflect changes in membrane architecture, surface antigen or host-protein display, cargo accessibility, or molecular organization. Complementary perturbation approaches will be required in the future to define the specific features of intact sEVs that promote efficient mucosal T cell priming.

## MATERIALS AND METHODS

### Bacterial culture and macrophage infection

*Salmonella enterica* serovar Typhimurium UK-1 was grown overnight in LB Miller broth at 37 °C with shaking at 250 rpm. Overnight cultures were diluted to an OD_600_ of 0.05 in 20 mL LB Miller broth and grown at 37°C with shaking to an OD_600_ of 0.5. Bacteria were then used to infect RAW264.7 macrophages at a multiplicity of infection (MOI) of 5. RAW264.7 macrophages were cultured to confluency in complete DMEM containing 10% fetal bovine serum (FBS) and 1% penicillin–streptomycin. Before infection, cells were washed with sterile phosphate-buffered saline, and the culture medium was replaced with antibiotic-and serum-free DMEM. Bacteria were collected by centrifugation, washed twice in sterile phosphate-buffered saline (PBS), and resuspended in PBS before adding to macrophage cultures. After 2 h of infection, medium was replaced with DMEM containing 1% exosome-depleted FBS and 100 μg/mL gentamicin for 1 hour. Cell cultures were then maintained in DMEM containing 1% exosome-depleted FBS and 25 μg/mL gentamicin until conditioned media were collected at 24 and 48 hours post-infection for downstream sEV isolation.

### sEV Isolation and Characterization

Conditioned media collected at 24 and 48 h post-infection were subjected to sequential centrifugation to remove cells, debris, and larger vesicles. Media were first centrifuged at 500 × g for 10 min, transferred to a new conical tube, centrifuged at 4,000 × g for 10 min, and then centrifuged at 16,000 × g for 30 min to remove cell debris and large EVs. The clarified cell culture supernatant was then passed through a 0.2 μm PES vacuum filter and subjected through two rounds of ultracentrifugation at 100,000 × g. The sEV pellets were collected in PBS supplemented with 1% protease inhibitor (Pierce Protease Inhibitor Tablets, Thermo Fisher, A32965). Isolated sEVs were characterized using ZetaView Quatt (Particle Metrix), used to determine particle size and concentration.

### Evaluation of vesicle integrity after sonication

Following sEV isolation and initial characterization, equal amounts of sEVs were aliquoted into two separate microcentrifuge tubes. One aliquot was disrupted using a probe sonicator for four cycles of 20 s ON and 60 s OFF, whereas the second aliquot was left intact. Samples were diluted with an equal volume of sterile phosphate-buffered saline (PBS) and loaded onto 2 mL 100 kDa molecular-weight cutoff (MWCO) Amicon centrifugal filter units (MilliporeSigma, UFC201024). Filters were centrifuged at 4,200 × g for 5 min to collect the flowthrough. Filter units were then inverted and centrifuged at 1,000 × g for 2 min to recover the retained fraction containing sEVs. Protein concentrations in the retained and flowthrough fractions from intact and disrupted sEV preparations were determined using a BCA assay (Thermo Fisher Scientific, 23227).

Vesicle morphology was assessed by transmission electron microscopy (TEM). Intact and sonication-disrupted sEV preparations were adsorbed onto electron microscopy grids, negatively stained, and imaged by TEM to evaluate vesicle morphology and membrane integrity.

Surface epitope preservation after sonication was assessed using the MACSPlex EV Kit IO (Miltenyi Biotec, 130-122-211) according to the manufacturer’s protocol. For each sample, 10 μg total EV protein was diluted in MACSPlex buffer and allocated to intact or disrupted sEV groups, with buffer alone included as a blank control. Samples were incubated overnight with MACSPlex EV IO Capture Beads, followed by washing and centrifugation. A 1:1:1 cocktail of EV IO Detection Reagents CD9, CD63, and CD81 (5 μL each) was added to each sample, followed by incubation and washing steps. Capture beads were resuspended, transferred to a 96-well round-bottom plate, and analyzed by flow cytometry. Data were analyzed according to the manufacturer’s instructions.

### Mouse Experiments

7-week-old female BALB/cJ mice (Jackson Laboratories, Strain# 000651) were used for in vivo immunization studies. For intranasal sEV immunization, mice were anesthetized using isoflurane and administered 40 μg sEVs per dose in 25 μL phosphate-buffered saline (PBS) containing 1% protease inhibitor cocktail. Mice received three total intranasal doses of intact sEV(+), sonication-disrupted sEV(+) (without Amicon filtration to normalize dose concentration), or PBS.

For live attenuated vaccine control immunization, mice first received 0.3 M sodium bicarbonate using a 22-gauge oral gavage needle and were allowed to rest for 10 min to neutralize stomach acid. Mice were then orally administered 5 x 10^9^ colony-forming units (CFUs) of ΔaroA *Salmonella* Typhimurium UK-1 (*18*), resuspended in 100 μL sterile phosphate-buffered saline using a clean gavage needle.

Seven weeks post first dose, mice were challenged orally with a lethal dose (4.5 x 10^6^ CFUs) of *Salmonella* Typhimurium UK-1 and euthanized 24 hours post challenge. For euthanasia, carbon dioxide was used as a primary method, followed by cervical dislocation as a secondary method. Blood samples were collected throughout the study in weeks 1, 3, 5, and 7 from the saphenous vein. Stool samples were collected at week 7, 24 hours post *Salmonella* challenge. Tissues harvested after euthanasia were dissociated using sterile syringes in PBS and resuspended in RPMI medium containing FBS and penicillin–streptomycin at 1 × 10⁶ cells per well in 6-well plates for ex vivo stimulation.

For the survival study, mice immunized with intact sEV(+), dis sEV(+), or PBS were challenged orally with a lethal dose (4.5 x 10^6^ CFUs) of *Salmonella* at week 7 post-immunization. Mice were followed till endpoint and weighed and body-scored daily in addition to assessment of dehydration. Mice were humanely euthanized upon reaching endpoint as determined by body scores and a loss of 20% baseline body weight. Log rank Mantel-Cox test was used to determine statistical significance.

### Staining of IgA-Coated Fecal Samples

Fecal pellets were collected from immunized mice one day post *Salmonella* challenge. The pellets were weighed and homogenized in sterile PBS on a vortex with shaker attachment for 15 minutes, followed by centrifugation at 800 x g for 5 minutes. The supernatant was collected in a separate tube and centrifuged at 9200 x g for 10 minutes to pellet the bacteria (*20*). Supernatant was discarded and bacterial pellet was resuspended in FACS buffer (PBS with 2% FBS) and centrifuged again. Bacterial cells were stained with FITC *Salmonella* antibody (Invitrogen, PA1-73020) and APC IgA antibody (Invitrogen, 17-4204-82) for 25 minutes in the dark at 4°C. Cells were washed with FACS buffer and resuspended in 250 μL FACS buffer to run through the flow cytometer.

### Opsonophagocytic Assay

Serum from week 7 post-immunization with either sEVs or saline was heat inactivated for 30 minutes at 56°C at a 1:50 dilution in PBS to get rid of complement factors. RAW264.7 macrophages were grown to confluency and counted. *Salmonella* Typhimurium UK-1 culture was grown up to 0.5 OD600 and volume needed for an MOI of 10:1 infection was collected and resuspended in 100 μL PBS. The calculated bacteria were mixed with the heat-inactivated serum and allowed to opsonize in a 37°C incubator for 30 minutes. Macrophages were infected with opsonized as well as un-opsonized bacteria for 30 minutes after which the media containing bacteria was removed and fresh media containing 100 μg/mL gentamicin was added for 1 hour to remove extracellular bacteria. Following this, media was removed, and cells were lysed using 0.1% Triton-X 100 for 15 minutes. Lysed cells were plated on agar plates which were incubated at 37°C overnight after which colonies were counted.

### Recombinant Protein Purification

The *Salmonella* Typhimurium UK-1 sequence was used to design primers for *sopB*: forward primer CAGGATCCGATGCAAATACAGAGCTTCTATCACTCAG, reverse primer GCTTGTCGATCAAGATGTGATTAATGAAGAAATGCCTTTTACT and *cirA*: forward primer TAAGCACATATGATGTTTAGGTTTAACCCTTTCGTTCGG, reverse primer TAAGCAGGATCCTTGAAACGGTAATCCACCGCCA. After running PCR, *sopB* and *cirA* gene amplification was confirmed using gel electrophoresis. The samples were cleaned using a PCR clean-up kit and cut using NdeI and BamHI restriction enzymes for 1 hour and 37°C. pACYCduet and pET28a vectors were used for *sopB* and *cirA* respectively. Digested vectors and gene samples were run on a gel and cut, followed by gel digestion to prepare the samples for ligation. Overnight ligation at 16°C was performed and bacterial transformation into *E. coli* BL21(DE3) was conducted by placing samples on ice for 30 minutes, a heat shock at 42°C for 2 minutes, followed by bacterial growth for 2 hours at 37°C with shaking at 250 rpm before plating the bacteria on chloramphenicol-and kanamycin-resistant LB agar plates. Individual transformed colonies were collected and grown overnight in LB miller. Plasmids were collected from colonies and confirmed through sequencing.

For recombinant protein expression, bacterial cultures were grown to an OD600 of 0.6, and protein expression was induced with 0.5 mM isopropyl β-D-1-thiogalactopyranoside (IPTG). Following this, bacterial cultures were grown overnight at 19°C with shaking at 250 rpm. Bacterial pellets were collected by centrifugation and resuspended in wash buffer containing 20 mM Tris-HCl, 200 mM NaCl, 5 mM imidazole, and 6 M urea. Cells were lysed using a probe sonicator for five cycles of 30 s ON and 60 s OFF. After this the buffer containing lysed bacterial cells was centrifuged again for 10 minutes at 4100 rpm. Supernatant was discarded and pellet was resuspended in 6 M urea solution and allowed to sit at room temperature for 10 minutes. The solution was centrifuged for 20 minutes at 4100 rpm after which the supernatant containing protein of interest was saved and passed through an equilibrated purification column containing a Ni-NTA resin (Thermo Scientific, 88221). The column was washed using wash buffer before and after the His-tagged protein was passed through it, after which it was eluted using an elution buffer (20mM Tris-HCl, 200mM NaCl, 200mM imidazole, and 6M urea). The collected samples were run on an SDS-PAGE and visualized using GelCode Blue (Thermo Scientific, 24590).

### Antibody Titer analysis by ELISA

Serum was isolated from blood upon centrifugation at 10,000 x g for 10 minutes and stored at-20°C until used for ELISAs. ELISA MaxiSorp 96-well plates (Thermo Scientific, 12-565-136) were coated with 2 μg recombinant antigen (OmpA, CirA, SopB) per well and the plate was stored overnight at room temperature. The unbound antigen was removed from the plate by blotting on a paper towel, and 200 μL per well blocking buffer (1% BSA in PBS) was added after which the plate was incubated at 37°C for 2 hours. After blocking, the wells were washed 3 times with PBST (PBS with 1% Tween-20) and 2 times with PBS and blotted dry each time. Serially diluted sera in ELISA buffer (PBS with 0.5% BSA and 0.05% Tween20) were added to the wells, and the plate was left at 4°C overnight. The plate was washed again with PBST and PBS after which the secondary goat anti-mouse IgG HRP antibody (Southern Biotech, 1030-05), goat anti-mouse IgG1 HRP (Invitrogen, A10551), or goat anti-mouse IgG2a HRP (Invitrogen, M32207) was added to the wells at a 1:2000 dilution in ELISA buffer, and the plate was incubated at 37°C for 90 minutes. After washing the plate again, 50 μL of pre-warmed ABTS substrate (2,2’-azino-bis(3-ethylbenzothiazoline-6-sulfonic acid)) was added and the plate was left at room temperature in the dark. The plate was run on the Cytation 3 plate reader (BioTek) at 415 nm and antibody log2 titers were calculated at an absorbance of 0.1U above negative control.

### Single-cell RNA Sequencing

Mesenteric lymph nodes were collected from sEV and PBS-immunized mice at week 7, 24 hours post *Salmonella* challenge. Sterile syringe plungers were used to make a single cell suspension of mLNs in sterile 1X PBS and cells were counted. Dead cell removal kit (Miltenyi, 130-090-101) was used to isolate live cells, after which cells were counted again and resuspended in 100 μL 1X PBS at a concentration of 2000 cells per μL. Samples underwent library prep using the 10x Chromium X and Single Cell 3′ Library & Gel Bead Kit v3 (10x Genomics) following manufacturers guidelines and total reads were assessed before proceeding to full sequencing. Approximately 20,000 single cells at 90% or greater viability were loaded into the Chromium Controller. Libraries were sequenced on the Illumina Novaseq X aiming for 50,000 reads per cell. Fastq files received after sequencing were uploaded to the 10X Genomics Cell Ranger cloud analysis software and the filtered and barcoded output files that were run against the mouse reference genome GRCm39 were used for downstream analysis using Seurat v5.4.0 on *R* Studio v4.5.2. Initial quality control was performed to filter cells based on mitochondrial content and number of genes detected, after which technical replicates were merged to identify clusters and create Uniform Manifold Approximation and Projection (UMAP) plots. Differentially expressed genes were identified, using FindAllMarkers on Seurat, between sEV-immunized and PBS control groups within selected immune cell compartments, including CD4 T cell clusters, excluding regulatory T cells, and separately for B cell clusters. Volcano plots displayed average log2 fold-change and −log10-adjusted p values, with genes classified as upregulated or downregulated using an absolute log2 fold-change threshold of 1 and an adjusted p value threshold of 0.05. Selected upregulated T cell and B cell genes were then grouped by enriched gene ontology categories and visualized as summary dot plots using *tidyverse* and *ggplot2*.

### *Ex vivo* Stimulation and Flow Cytometry

Single-cell suspensions from mesenteric lymph nodes (mLNs) and axillary lymph nodes (axLNs) were plated in 6-well plates at 1 × 10⁶ cells per well. Cells were stimulated *ex vivo* for 72 hours with 15 μg sEV protein from uninfected macrophages [sEV(−)], 15 μg sEV protein from *Salmonella*-infected macrophages [sEV(+)], 15 μg disrupted sEV protein from *Salmonella*-infected macrophages [disrupted sEV(+)], a recombinant antigen cocktail containing OmpA, SopB, OmpD, FliC, and CirA at 1 μg per antigen, or left unstimulated. Protein transport inhibitor cocktail (Thermo Fisher Scientific, 00-4980-93) was added during the final 5 h of stimulation. Cells were then collected in 5 mL FACS tubes and centrifuged at 500 × g for 5 min. Cell pellets were washed twice with PBS, stained with Zombie Green viability dye for 10 min at 4°C, and transferred to a 96-well plate. Samples were washed twice with FACS buffer and blocked with Fc block for 5 min at room temperature in the dark. Cells were then surface stained for 25 min at 4 °C with an antibody cocktail for CD4 T cells (Panel 1) containing BV421-CD45, BV605-TCRb, APC/Cy7-CD4, PE/Cy5-CD62L, and APC-CD44, and another for CD8 T cells (Panel 2) containing BV421-CD8, BV605-TCRb, APC/Cy7-CD4, PE/Cy5-CD62L, and APC-CD44 (BioLegend). Samples were washed twice with FACS buffer at 500 × g for 5 min and fixed/permeabilized using Cytofix/Cytoperm solution (BD Biosciences) for 35 min. Upon fixation, intracellular cytokine staining was conducted wherein cells were washed twice with 1X permeabilization buffer (BD Biosciences), followed by staining with AF700-IL2 and PE-IFNγ for 45 minutes at 4°C. Cells were washed again with 1X permeabilization buffer. Finally, cells were resuspended in 200 μL FACS buffer and run on a CytoFLEX flow cytometer, followed by analysis using FlowJo v10 (Beckman Coulter).

### Primary Bone Marrow Derived Dendritic Cells (BMDC) Isolation and Differentiation

7–8-week-old male BALB/cJ (Jackson Laboratories, Strain# 000651) mice were housed under specific-pathogen free conditions at the University of Florida’s animal facility. Bone marrow was collected from mice and cells were cultured in Iscove’s Modified Dulbecco’s Medium (IMDM) containing 10% FBS, 1% penicillin/streptomycin, Gentamycin (50 µg/mL), 1X GlutaMax, Betamercaptoethanol (55 µM), and Granulocyte-Macrophage Colony-Stimulating Factor (GM-CSF) (3 ng/mL; BD-Bioscience 554586). BMDCs are a mixed population of adherent and semi-adherent cells, so additional care was taken to retain cell density. Flow cytometry was utilized to confirm the differentiation status and surface marker expression of the BMDC population.

### BMDC *ex vivo* sEV Stimulation and CD4 T Cell Co-culture

6-well cell culture plates were seeded with isolated BMDCs at 1×10^7^ cells/well in complete IMDM. On Day 2, 1 mL of complete IMDM was added to each well. On Day 4, 1 mL of cell culture medium was removed from each well and replaced with 1 mL of complete IMDM. On Day 6 (and each day on until sEV stimulation), 1 mL of cell culture medium was removed from each well and replaced with 1 mL of complete IMDM. 2 μg sEVs(+) or dis sEVs(+) were added to wells in triplicate, and additional wells were left unstimulated or used as count wells. BMDCs were collected after 24 hours of sEV stimulation and stained for flow cytometry with BV421-CD3+CD19 (dump gate), PE-MHCII, APC-CD11c, AF700-CD11b, Zombie Green Live/Dead, BV605-CD86, PE/Cy5-CD80 (Biolegend). Samples were run on a flow cytometer (CytoFLEX, Beckman Coulter) and analyzed using FlowJo v10.

Additional wells containing BMDCs stimulated with intact and dis sEV(+) for 24 hours received naïve CD4 T cells at a ratio of 5:1 (T cells:BMDCs) isolated from a mixture of spleens, mesenteric, inguinal, and axillary lymph nodes using a negative CD4 T cell isolation kit (StemCell, 19812A). T cells were allowed to activate for 48 hours after which cells were collected and stained for flow cytometry with Zombie Green Live/Dead, BV421-CD3, PE/Cy5-CD69, APC/Cy7-CD4, PE-IFNγ, AF700-IL2 (Biolegend), and analyzed using FlowJo v10.

## Statistical analysis

Statistical analyses were performed using GraphPad Prism version 11, except for single-cell RNA-sequencing analyses, which were performed in R as described above. Biological replicate numbers (*n*) are indicated in the corresponding figure legends. Independent mice or independently prepared sEV isolations were considered biological replicates, as specified for each experiment. Technical replicates were not treated as independent biological replicates. All statistical tests were two-sided, and *P* < 0.05 was considered statistically significant.

## Acknowledgments

We thank Dr. Roy Curtiss III for providing the *Salmonella* strains used in this study and Dr. Rhonda Bacher for helpful discussions. Transmission electron microscopy experiments were performed at the UF ICBR Electron Microscopy core (RRID:SCR_019146). We also thank Brandon Wilson and Dr. Yanping Zhang at the University of Florida Interdisciplinary Center for Biotechnology Research (UF ICBR) for assistance with single-cell RNA sequencing. Single-cell library preparation was performed by the UF ICBR Gene Expression & Genotyping Core (RRID:SCR_019145), and sequencing was performed by the UF ICBR NextGen DNA Sequencing Core (RRID:SCR_019152).

## Author contributions

Conceptualization: SB, MJF, Methodology: SB, ACS, ST, MJF, Investigation: SB, ACS, ST, Visualization: SB, MJF, Funding acquisition: MJF, ACS, Project administration: MJF, Supervision: MJF, Writing – original draft: SB, ACS, ST, MJF

## Data and materials availability

The single-cell RNA-seq raw sequencing data in this study have been deposited in the Gene Expression Omnibus under accession number GSE336745. Processed files include raw and filtered gene-barcode matrices, cell-level metadata, cluster annotations, UMAP coordinates, and differential expression tables. Other scRNA-seq data are included as supplementary tables.

## Ethics Statement

All animal procedures were approved by the University of Florida Institutional Animal Care and Use Committee (IACUC 202200000015) and performed in accordance with institutional and federal guidelines.

