## Supplemental data for "Vesicle architecture licenses antigen-specific memory T cell immunity after mucosal immunization with infection-derived vesicles"

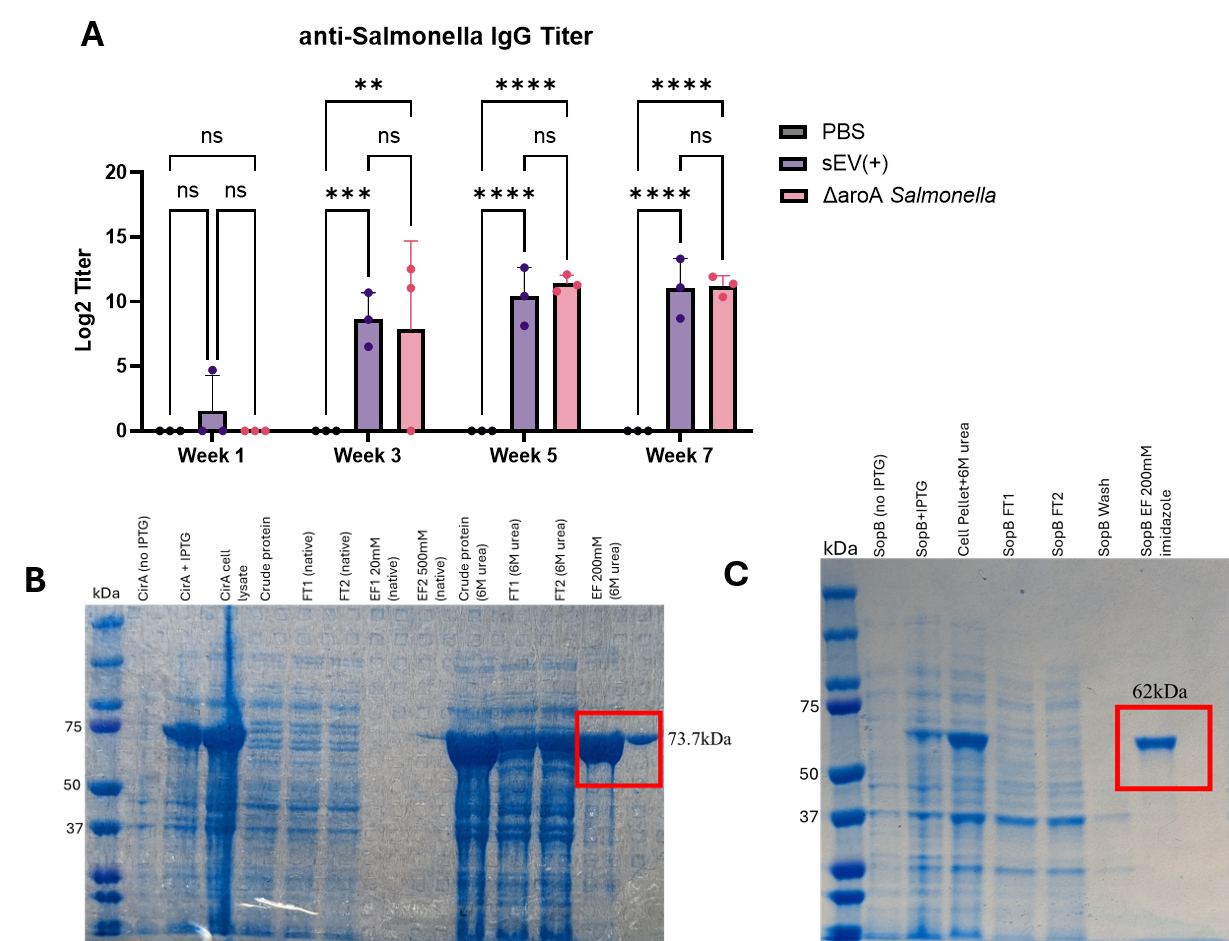

**Figure S1: sEV(+) immunization leads to increasing IgG production against whole *Salmonella* lysate. (A)** Log2 titers of *Salmonella-*specific serum IgG levels in mice immunized with PBS, sEV(+), or ΔaroA *Salmonella* over the span of seven weeks*.* Two-way ANOVA was used to determine statistical significance, shown as ** (p<0.01), *** (p<0.001), and **** (p<0.0001). **(B-C)** SDS-PAGE showing His-tagged purified recombinant *Salmonella* CirA and SopB proteins, purified using affinity chromatography.

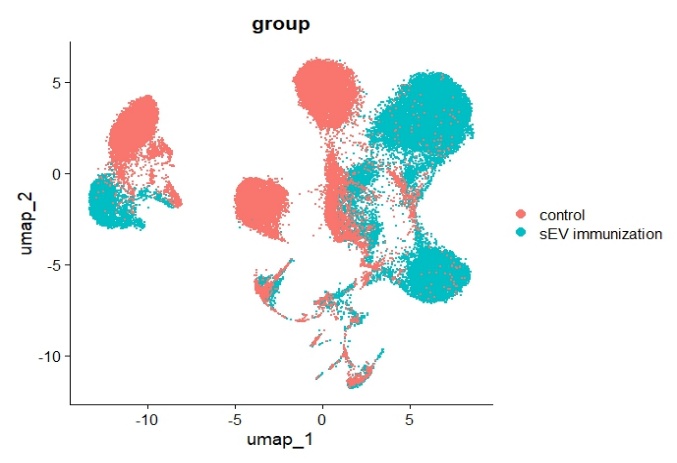

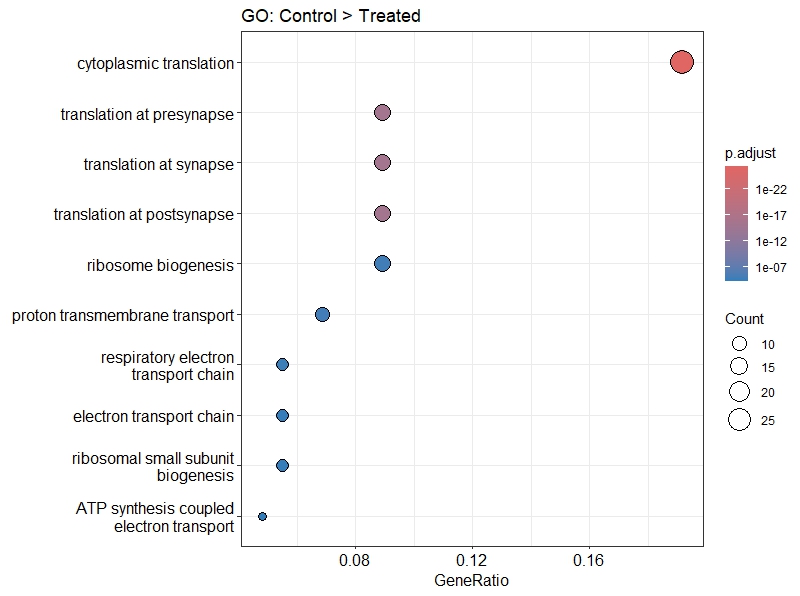

**A**

**B**

**C**

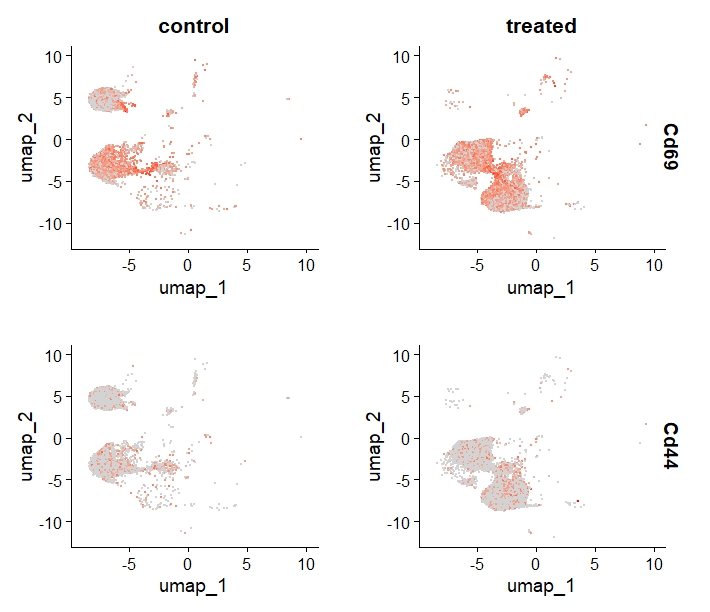

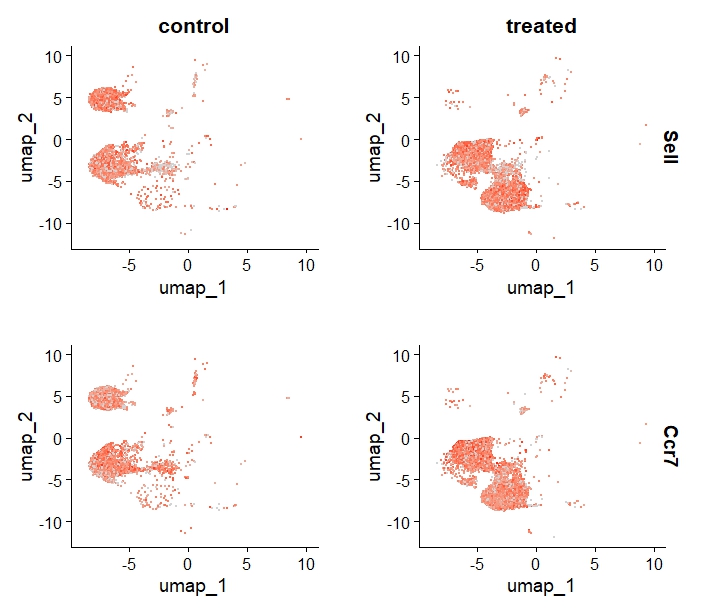

**E**

**D**

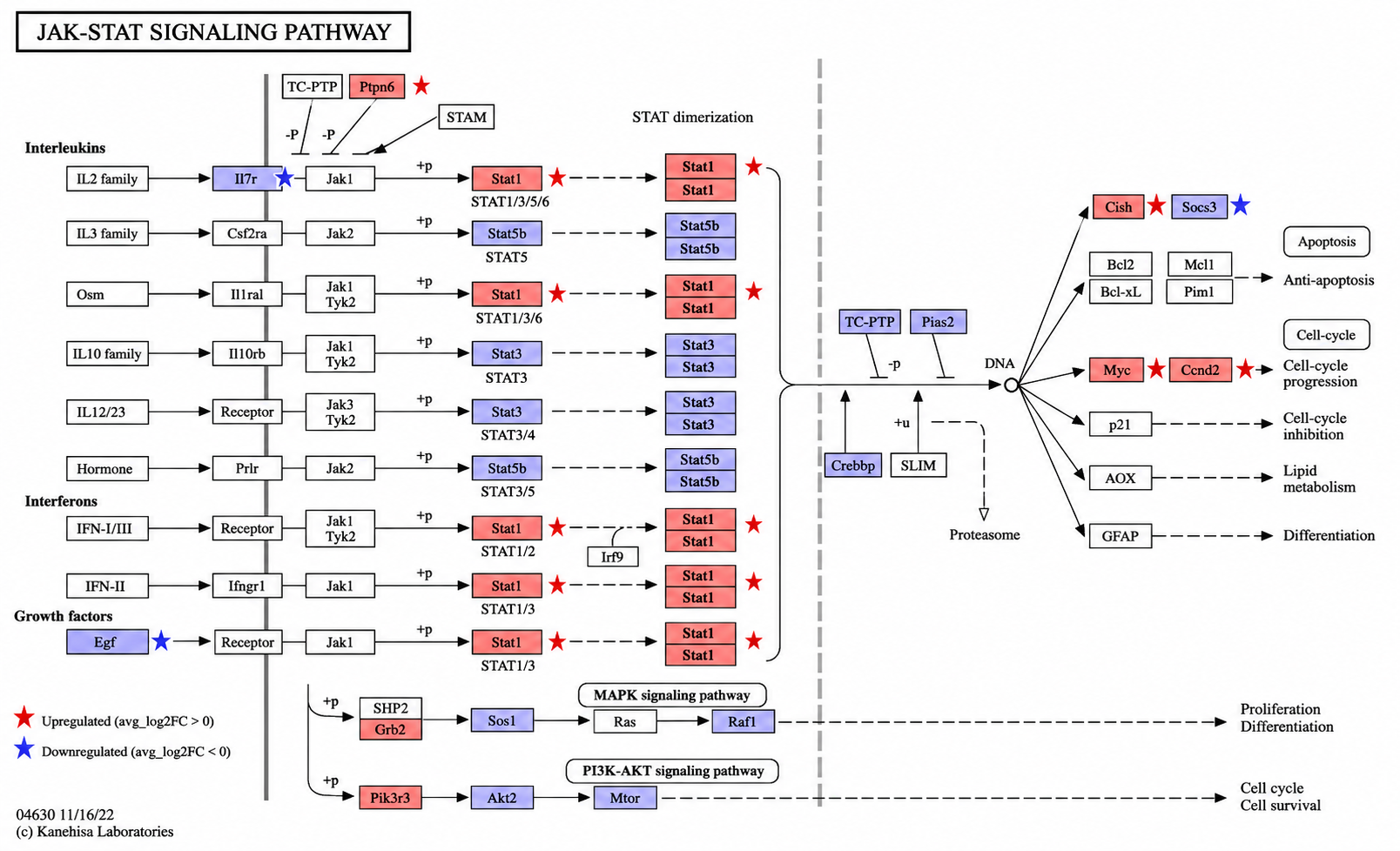

**Figure S2: sEV(+) immunization promotes T cell activation and interferon-associated transcriptional programs after lethal *Salmonella* challenge. (A)** UMAP visualization of T cell clusters from PBS control and sEV(+)-immunized mice after lethal *Salmonella* challenge. **(B)-C)** Feature plots showing expression of CD69, CD44, Sell, and CCR7 in control and sEV(+)-immunized T cell clusters. **(D)** KEGG pathway mapping of differentially expressed genes onto the JAK–STAT signaling pathway in T cell clusters from sEV(+)-immunized mice compared with PBS controls. Red indicates increased expression and blue indicates reduced expression in sEV(+)-immunized mice. Stars show significant change. **(E)**  Gene ontology enrichment analysis of pathways enriched in control CD4+ T cell clusters compared with sEV(+)-immunized CD4+ T cell clusters (excluding Tregs).

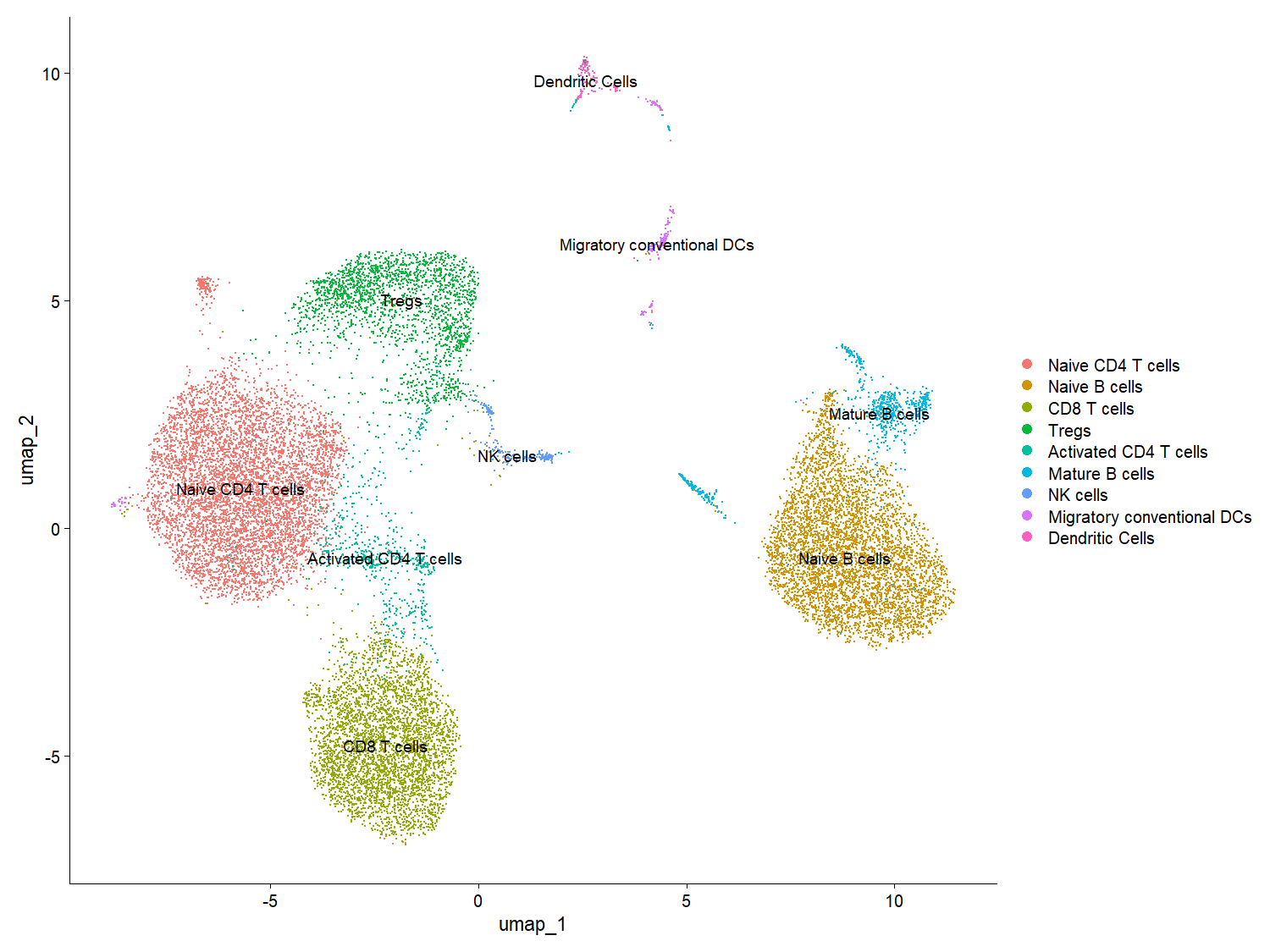

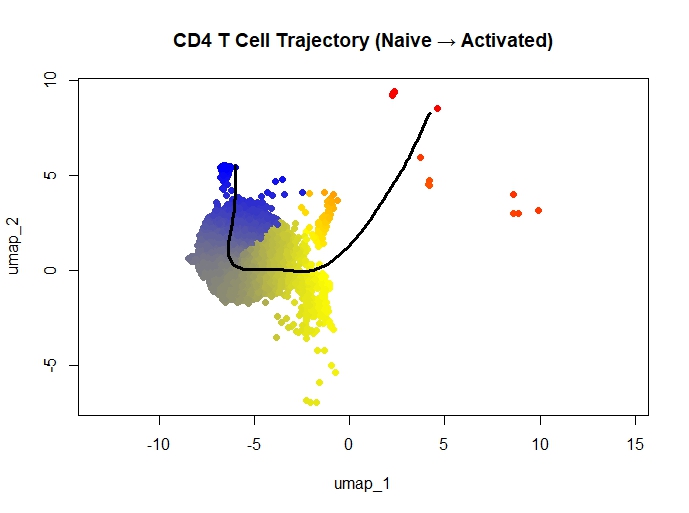

**A**

**B**

**Figure S3: Immunization with PBS does not lead to memory T cell activation upon *Salmonella* challenge. (A)** UMAP visualization of all identified clusters from PBS-immunized mice. **(B)** Pseudotime trajectory analysis, conducted using Slingshot on R studio, demonstrating predicted trajectory from naïve to activated CD4 T cell clusters as shown in **(A)** from mice immunized with PBS control.

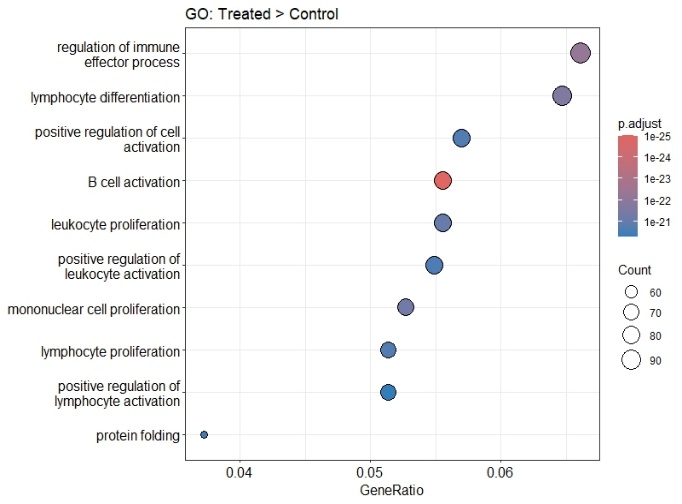

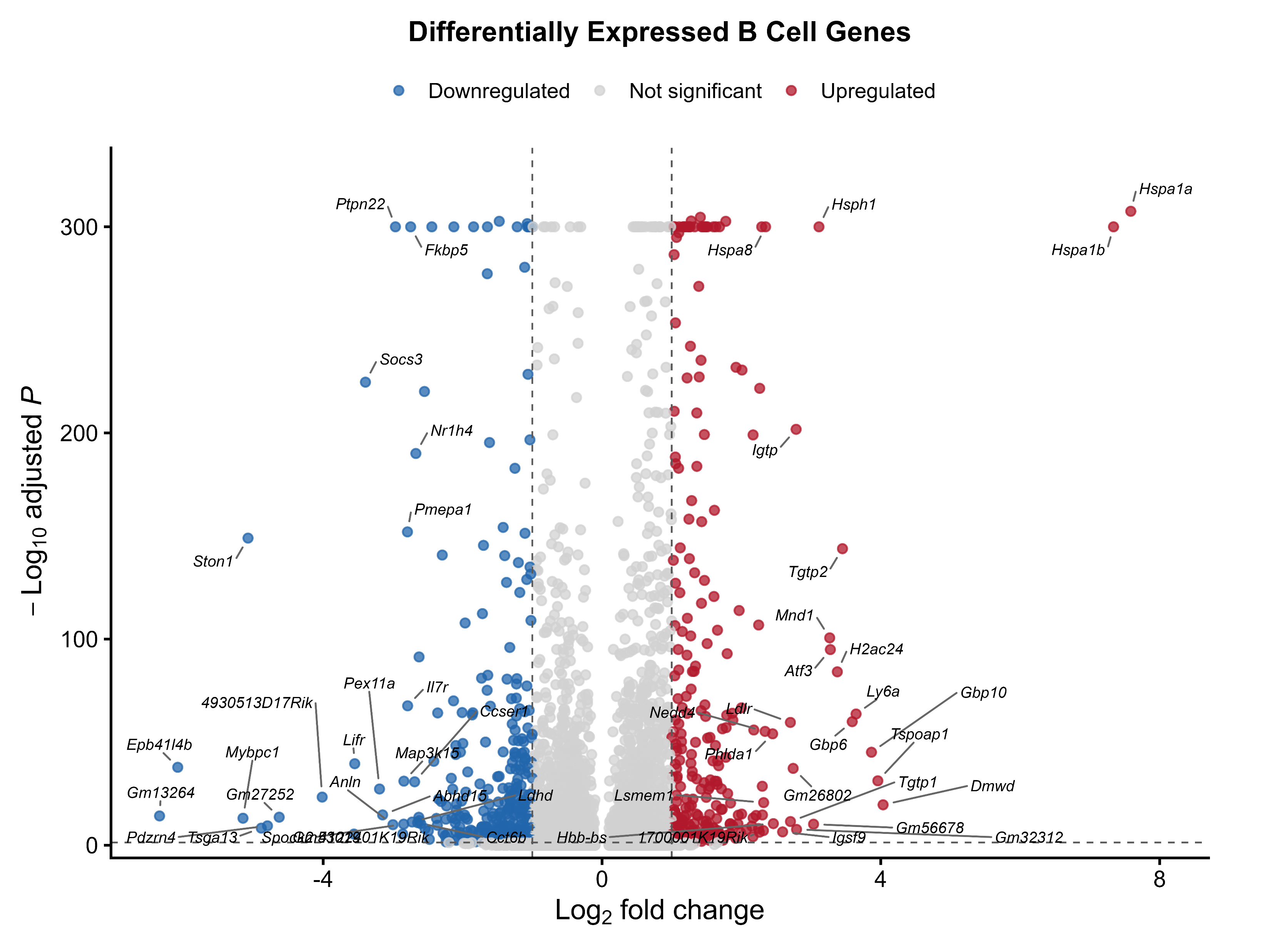

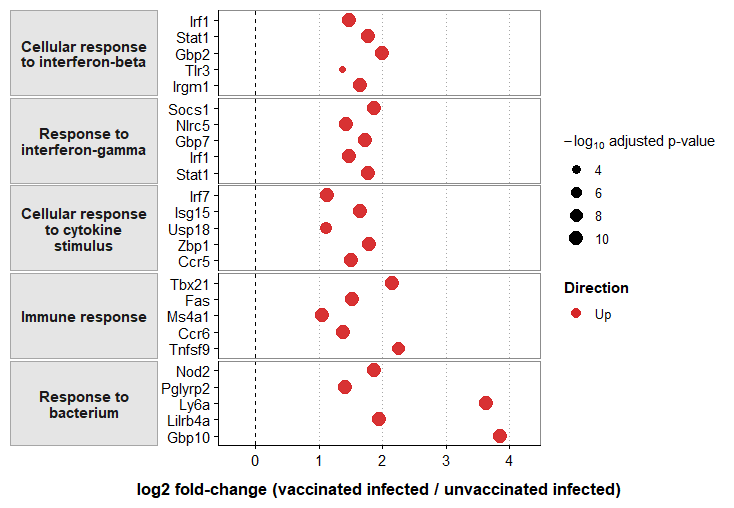

**log2FC sEV (+)/control**

**A**

**B**

**C**

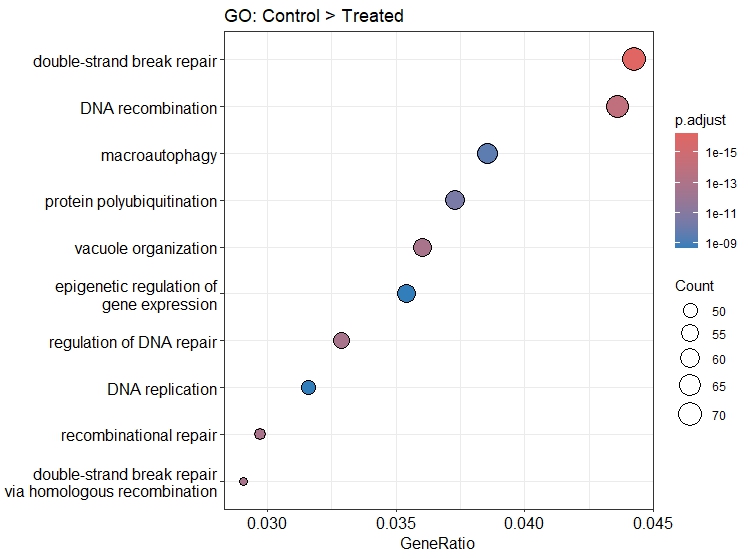

**D**

**Figure S4: sEV(+) immunization promotes B cell activation after lethal *Salmonella* challenge. (A)** GO enrichment pathway analysis of differentially expressed genes in all B cell clusters of treated [sEV(+)-immunized] versus control (PBS-immunized) mice conducted on R studio. **(B)** GO enrichment pathway analysis of differentially expressed genes in all B cell clusters of control (PBS-immunized) versus treated [sEV(+)-immunized] mice conducted on R studio. **(C)** Volcano plot showing the expression data across all B cell clusters. **(D)** Select gene expression analysis from **(C).**

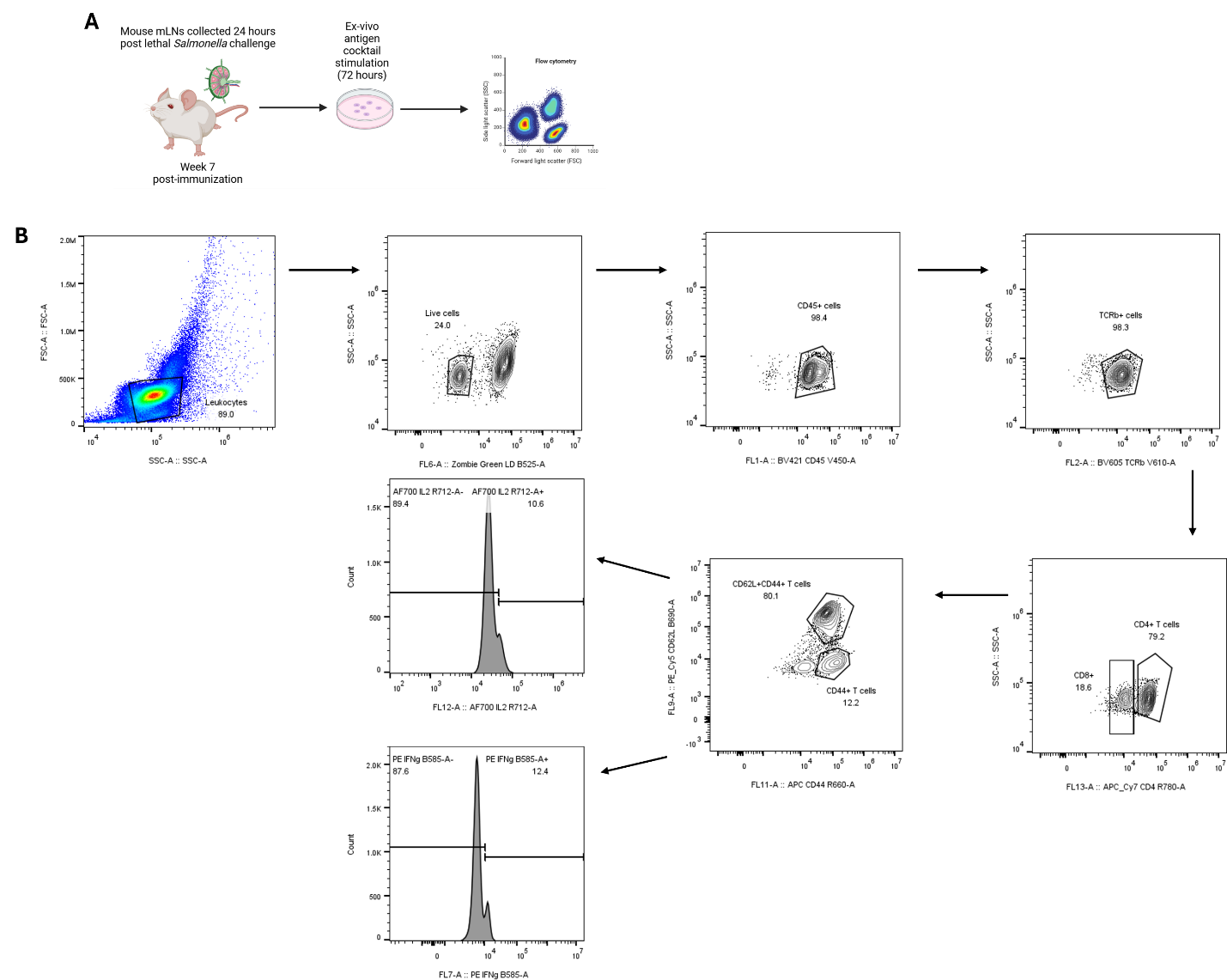

**Figure S5:** **Ex-vivo stimulation with *Salmonella* recombinant antigen cocktail leads to memory T cell activation. (A)** Biorender diagram showing ex-vivo stimulation scheme at week 7 post-immunization with intact *Salmonella*-infection derived macrophage sEVs [sEV(+)], disrupted sEVs from *Salmonella-*infected macrophages [dis sEV(+)], or PBS. Mesenteric lymph node (mLN) cells were stimulated for 72 hours with recombinant *Salmonella* antigen cocktail consisting of OmpA, OmpD, FliC, SopB, and CirA prior to staining for flow cytometry. **(B)** Flow cytometry gating strategy used to identify antigen-specific memory CD4 or CD8 T cells and their intracellular cytokine production.

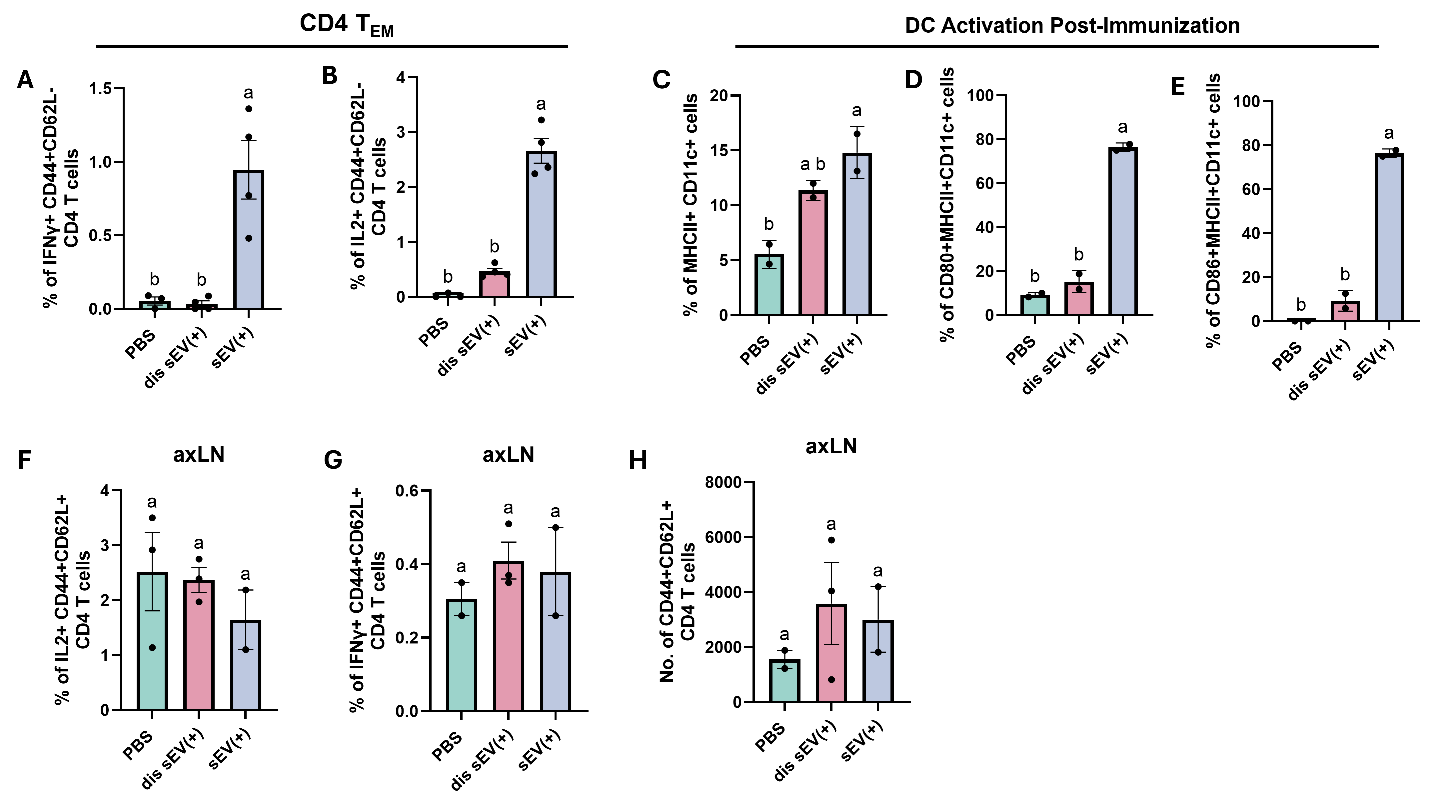

**Figure S6: Immunization with intact sEV(+) induces spatial activation of effector memory CD4 T cells and dendritic cells in the mesenteric lymph nodes (A-B)** Percentages of IFNγ- and IL-2-expressing CD4 T_EM_ cells (CD44+CD62L-) after ex vivo stimulation of mesenteric lymph node cells with recombinant *Salmonella* antigen cocktail (n=4 mice per immunization group). **(C)** Percentage of MHCII+CD11c+ DCs isolated from mLNs of *Salmonella*-challenged mice immunized with intact sEV(+), disrupted sEV(+), or PBS, at week 7 post-immunization. **(D)** Percentage of CD80+MHCII+CD11c+ DCs isolated from mLNs of *Salmonella*-challenged mice immunized with intact sEV(+), disrupted sEV(+), or PBS, at week 7 post-immunization. **(E)** Percentage of CD86+MHCII+CD11c+ DCs isolated from mLNs of *Salmonella*-challenged mice immunized with intact sEV(+), disrupted sEV(+), or PBS, at week 7 post-immunization (n=2 mice per immunization group). **(F-G)** Percentage of IFNγ- and IL2-expressing axillary lymph node central memory (T_CM_) CD4 T cells in mice immunized with intact sEV(+), disrupted sEV(+), or PBS. **(H)** Number of central memory (T_CM_) CD4 T cells in axillary lymph nodes of mice immunized with intact sEV(+), disrupted sEV(+), or PBS (n=2-3 mice per immunization group). Statistical significance was determined using one-way ANOVA with multiple comparisons, shown using compact letter display (a,b). Data shown as mean +/- SEM.

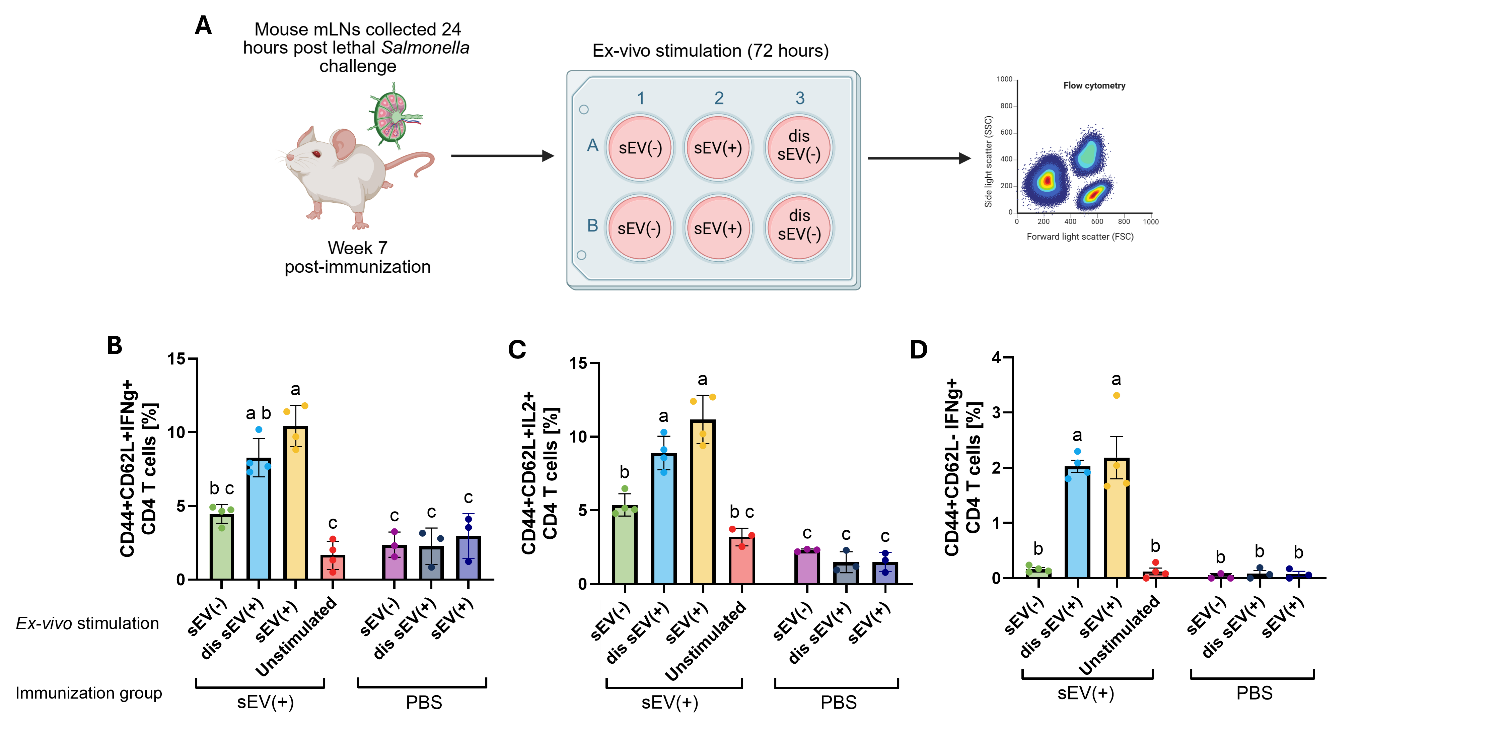

**Figure S7:** **Ex-vivo stimulation with disrupted and intact sEVs from *Salmonella-*infected macrophages leads to memory T cell activation. (A)** Biorender diagram showing ex-vivo stimulation scheme at week 7 post-immunization with sEV(+) or PBS. Mesenteric lymph node (mLN) cells were stimulated with sEVs from uninfected macrophages [sEV(-)], intact sEVs from infected macrophages [sEV(+)], disrupted sEVs from infected macrophages [dis sEV(+)] or left unstimulated prior to staining for flow cytometry. **(B)** Flow cytometry graph displaying percentage of IFNγ-expressing central memory CD4 T cells (CD4+CD44+CD62L+) from intact sEV(+)- or PBS-immunized mice in response to ex-vivo stimulation with sEVs from different conditions or no stimulation. **(C)** Flow cytometry graph displaying percentage of IL2-expressing central memory CD4 T cells (CD4+CD44+CD62L+) from intact sEV(+)- or PBS-immunized mice in response to ex-vivo stimulation with sEVs from different conditions or no stimulation. **(D)** Flow cytometry graph displaying percentage of IFNγ-expressing effector memory CD4 T cells (CD4+CD44+CD62L-) from intact sEV(+)- or PBS-immunized mice in response to ex-vivo stimulation with sEVs from different conditions or no stimulation. Statistical significance was determined using one-way ANOVA, shown as compact letter display (a,b,c).
